# A gene expression atlas of embryonic neurogenesis in *Drosophila* reveals complex spatiotemporal regulation of lncRNAs

**DOI:** 10.1101/483461

**Authors:** Alexandra L. McCorkindale, Philipp Wahle, Sascha Werner, Irwin Jungreis, Peter Menzel, Chinmay J. Shukla, Rúben Lopes Pereira Abreu, Rafael Irizarry, Irmtraud M. Meyer, Manolis Kellis, Robert P. Zinzen

## Abstract

We present a spatiotemporal transcriptome during early *Drosophila* embryonic nervous system development, revealing a complex cell type-specific network of mRNAs and lncRNAs.

**Abstract:** Cell type specification during early nervous system development in *Drosophila melanogaster* requires precise regulation of gene expression in time and space. Resolving the programs driving neurogenesis has been a major challenge owing to the complexity and rapidity with which distinct cell populations arise. To resolve the cell type-specific gene expression dynamics in early nervous system development, we have sequenced the transcriptomes of purified neurogenic cell types across consecutive time points covering critical events in neurogenesis. The resulting gene expression atlas comprises a detailed resource of global transcriptome dynamics that permits systematic analysis of how cells in the nervous system acquire distinct fates. We resolve known gene expression dynamics and uncover novel expression signatures for hundreds of genes among diverse neurogenic cell types, most of which remain unstudied. We also identified a set of conserved and tissue-specifically regulated long-noncoding RNAs (lncRNAs) that exhibit spatiotemporal expression during neurogenesis with exquisite specificity. LncRNA expression is highly dynamic and demarcates specific subpopulations within neurogenic cell types. Our spatiotemporal transcriptome atlas provides a comprehensive resource to investigate the function of coding genes and noncoding RNAs during critical stages of early neurogenesis.

## Introduction

Development of complex tissues from naïve primordia requires the precise spatiotemporal deployment of transcriptional programs as cells subdivide, specify, and differentiate. Owing to the availability of tissue- and cell type-specific markers characteristic for neurogenic cell types in the fruit fly embryo (Heckscher et al. 2014), *Drosophila* neurogenesis is highly tractable and several crucial regulators of neurogenesis have been identified over the past several decades (Skeath & Thor 2003; Beckervordersandforth et al. 2008; Broadus et al. 1995; Landgraf et al. 1997; Rickert et al. 2011; Wheeler et al. 2006; Doe 2017; Heckscher et al. 2014; Skeath et al. 1994; Weiss et al. 1998; Wheeler et al. 2009). Among the earliest events in embryonic neurogenesis is the subdivision of the lateral neurogenic ectoderm into columnar domains along the dorsoventral axis (Ohlen & Doe 2000a; Cowden & Levine 2003). This is followed by the formation of proneural clusters and consecutive phases of delamination, where neuroblasts cease contact with surrounding cells of the neuroectodermal columns and ingress into the embryo (Campos-Ortega 1995). Embryonic neuroblasts – *Drosophila* neural stem cells – undergo a series of self-renewing asymmetric divisions that produce ganglion mother cells, which give rise to glia and neurons (Broadus et al. 1995; Sousa-Nunes et al. 2010; Homem & Knoblich 2012; Heckscher et al. 2014). Importantly, each of the three neurogenic columns gives rise to molecularly and functionally distinct sets of neuroblasts (Doe 1992), but the molecular mechanisms that link spatial origin to the ensuing distinct fates remain poorly understood. To date, a small set of marker genes specifically expressed in individual columnar domains and in emerging cell types has been identified, but it remains unclear how these cell populations differ with respect to the global gene expression programs that shape their identities.

While expression dynamics of protein-coding transcripts have given important insights into the mechanisms that drive cellular differentiation, it should be noted that an emerging class of noncoding transcripts – the long noncoding RNAs (lncRNAs) – may well emerge as pivotal regulators of neurogenesis. In mammals, lncRNAs have been shown to be especially abundant in differentiated neuronal cells (Briggs et al. 2015), are expressed often with exquisite spatiotemporal specificity in the nervous system (Sauvageau et al. 2013; Goff et al. 2015), and some lncRNA species even exhibit neuronal subtype specificity (Molyneaux et al. 2015; S. J. Liu et al. 2016). Though the functional importance of some lncRNAs for development and cellular identity has been demonstrated in *Drosophila* (Wen et al. 2016), including in the nervous system ((Li & Liu 2015);(Landskron et al. 2018), very little is known about the cell type specific expression and function of lncRNAs over the course of early neurogenesis.

Large-scale efforts have characterized spatial gene expression in RNA *in situ* hybridization screens (Tomancak et al. 2002; Inagaki et al. 2005; Tomancak et al. 2007; Lécuyer et al. 2007; Wilk et al. 2016), but such efforts are qualitative rather than quantitative and largely exclude lncRNAs. In contrast, efforts to determine global transcriptome dynamics in the developing *Drosophila* embryo (Graveley et al. 2011; Brown et al. 2014; Young et al. 2012; B. Chen et al. 2016) may detect the expression of lncRNAs, but lack cell type resolution. As for most complex tissues, recapitulating early neurogenesis in cell culture is unfortunately not an option, because accurate specification and differentiation of cells depends on embryonic context, intricate interactions among cells within the neuroectoderm (Kunisch et al. 1994; Lai 2004) and signaling gradients involving surrounding tissues (Bier & De Robertis 2015; Rogers et al. 2017).

To overcome these limitations and to dissect stage- and cell type-specific transcriptomes in early neurogenesis, we adapted MARIS (Hrvatin et al. 2014) for use in developing *Drosophila* embryos. DIV-MARIS (***D**rosophila In **V**ivo* **M**ethod for **A**nalyzing **R**NA following Intracellular Sorting) allows purification of chemically cross-linked cell types from staged developing embryos based on marker gene expression, followed by RNA extraction and next-generation sequencing. Here, we employ DIV-MARIS to determine the transcriptome dynamics in distinct neurogenic cell populations. We assess the gene expression programs of two principal neurogenic domains – the ventral- and the intermediate columns, and of three differentiating cell types (neuroblasts, neurons and glia) at consecutive time points from primordial specification and subdivision to terminal differentiation.

DIV-MARIS reveals an extensive network of dynamic spatiotemporal gene expression during embryonic nervous system development. Our method reliably identifies known cell type-specific markers, but also reveals novel expression features. Furthermore, we uncover many genes – most of which have conserved homologs in human – that are expressed in distinct cell types throughout early neurogenesis and whose functions remain to be elucidated. Hence, DIV-MARIS provides an accurate expression map of spatiotemporal transcriptional programs driving early nervous system development. Moreover, our analyses identified many lncRNAs expressed in cell type-specific patterns and for which no functional roles are yet known. Applying stringent criteria for selection, we characterize 13 neural cell type-enriched lncRNAs with varied temporal expression, abundance, and subcellular localization. *In situ* visualization of lncRNA expression exposes an additional layer of specificity as neuroglial lncRNAs tend to be expressed highly, but only in extremely distinct subpopulations.

This study delivers a genome-wide, yet cell-type-specific view of gene expression during *Drosophila* neurogenesis from neurogenic columns to differentiated neurons and glia, provides insights into the expression properties of the coding and noncoding transcriptomes and will serve as a valuable tool for understanding how regulated coding and non-coding gene expression drives cell fate determination in early neurogenesis.

## Results

### Isolation of neuroglial cell types with spatiotemporal resolution

Early *Drosophila* neurogenesis starts with the specification of the lateral neurogenic ectoderm at the onset of zygotic transcription. The neurogenic ectoderm is quickly subdivided into distinct neurogenic columns (Ohlen & Doe 2000b; Cowden & Levine 2003), from which neuroblasts delaminate and undergo asymmetric division giving rise to ganglion mother cells (GMCs), followed by differentiation of GMCs into neurons and glia (Fig. 1A). To dissect the genome-wide transcriptional programs driving early neurogenesis, we purified specific cell populations comprising the neuroglial lineages using fluorescence-activated cell sorting (FACS) of chemically fixed cells. We isolated cells of the intermediate column (IC) and the ventral column (VC) using transgenic constructs by fusing IC-or VC-specific enhancers to reporter genes (Fig. S1A, 1B). Neuroblasts/GMCs, neurons, and glia cells were purified using antibodies directed against the endogenous markers *prospero (pros), embryonic lethal abnormal vision (elav), and reversed polarity (repo),* respectively (Fig. 1B, 1D, S1B). Early neurogenesis is a rapidly unfolding process, with naïve primordia developing into differentiated cell types in a matter of hours (Fig. S2A). To assess the temporal dynamics of early neurogenesis, we collected these cell populations at developmental stages (bins) that encompass critical events along the neurogenic lineages from early specification to terminal differentiation (Fig. 1B, S2A). Timed embryo collections were manually staged to assure which neurogenic events were captured within the collection bins (Fig. S2B). The earliest collection bin (4-6h after egg laying, AEL) primarily contains embryos immediately after specification and subdivision of the neurogenic ectoderm and encompasses the first rounds of neuroblast delamination. The second bin (6-8h AEL) includes all waves of neuroblast delamination, proliferation and diversification, followed by early differentiation in the third bin (8-10h AEL). A later collection towards the end of embryogenesis (18-22h AEL) serves as a reference point for fully differentiated neurons and glia.

**Fig. 1.**
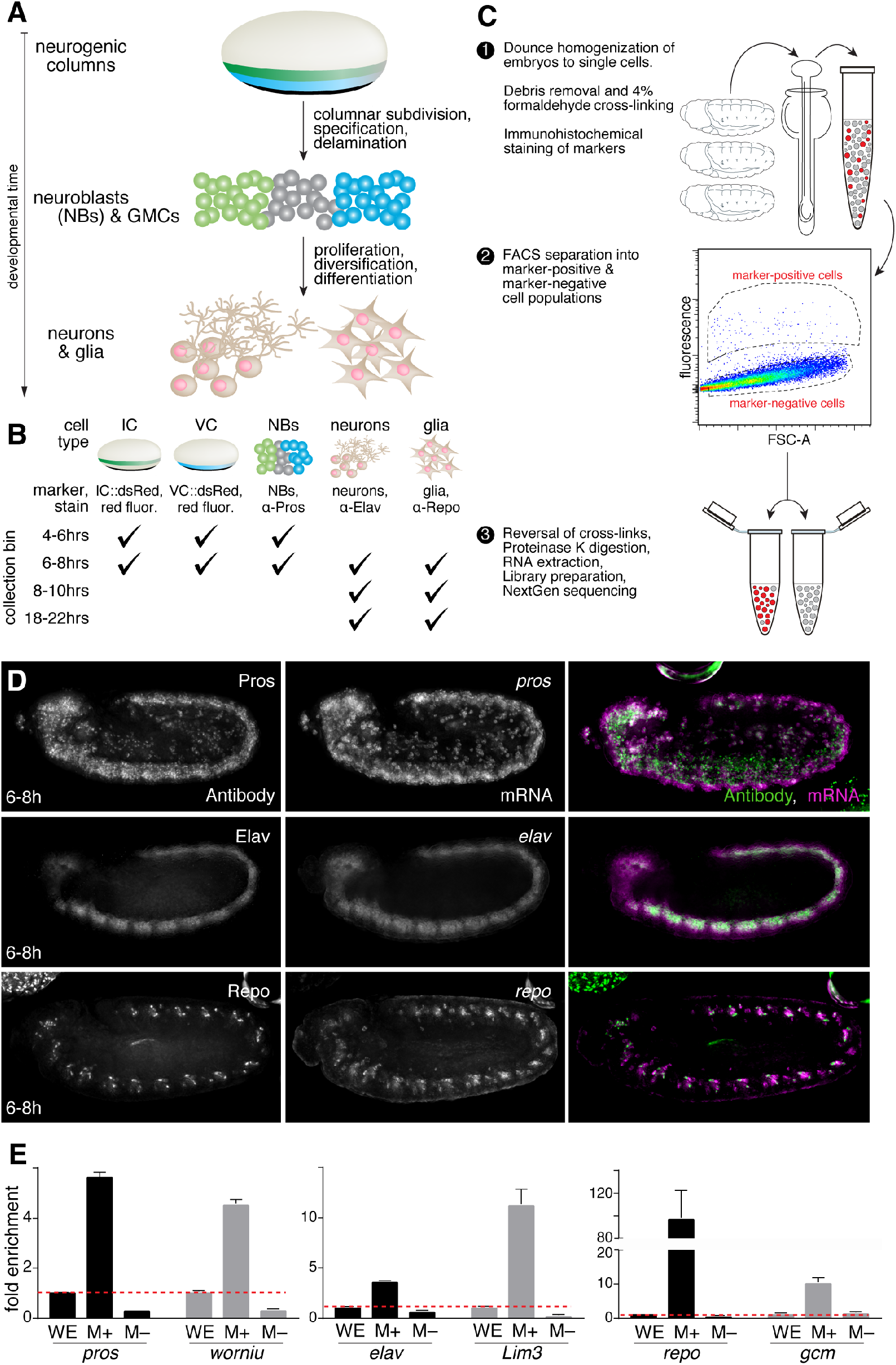
*DIV*-MARIS for the enrichment of staged neurogenic cell types. (A) Biological material enriched over the course of neurogenesis, including intermediate column (IC, green), ventral column (VC, cyan), neuroblasts (NBs), neurons, and glia. (B) Time windows of embryonic development (“collection bins”) targeted in this study, and the cell types isolated for each. (C) Schematic overview of *DIV-MARIS* protocol. (D) Multiplex whole mount immunohistochemistry (left) and RNA-FISH (middle) show faithfulness of the antibody sorting markers for neuroblasts (top row, Pros=prospero), neurons (middle row, Elav), and (F) glia (bottom row, Repo) at 6-8h. (E) Expression of marker genes used for FACS (black bars) and additional marker genes specific to the cell type of interest (grey bars) as measured by qPCR in marker-enriched (M+), and marker-depleted (M-) populations, calculated relative to whole embryo (WE, dashed red line); embryos collected at 4-10h AEL (*n* = 2). Error bars represent mean+s.e.m. C_T_ values normalized to the mean of two ubiquitous reference genes, *Actin42A* and *α-Tubulin.*

To isolate cell type-specific RNA from specific neurogenic cell types we adapted the MARIS protocol (Hrvatin et al. 2014), but had to introduce several modifications to temporally resolve cell types from complex and quickly developing *Drosophila* embryos *in vivo.* DIV-MARIS (outlined in Fig. 1C) is a flexible method for the isolation of high-quality RNA from specific fixed cell types within complex and rapidly developing embryos. Briefly, staged embryos are collected, dissociated into single-cell suspensions, and immediately cross-linked with formaldehyde. Neurogenic cell types were stained using antibodies, either against transgenic reporters (for the ventral and intermediate columns, Fig. S2A), or against endogenous markers (for neuroblasts/GMCs, neurons, and glia; Fig. 1D, Fig. S2B). Positively marked and unmarked populations were purified by FACS (Fig. 1C). We ascertained that the sorting strategy reliably isolated marked cells of interest by microscopy (e.g. Fig. S1C), as well as by analytical cytometry (e.g. Fig. S3); samples generally had purities >95% and samples below 90% purity were discarded. Furthermore, we evaluated the enrichment of DIV-MARIS-sorted cell types by quantitative RT-PCR against several marker genes associated with the cell types of interest (i.e. *pros* & *worniu* in neuroblasts/GMCs, *elav* & *Lim3* in neurons, *repo* & *gcm* in glia) as independent measures of cell type enrichment (Fig. 1E). We confirmed specific enrichment of the expected markers in sorted cells compared with whole embryos, as well as their depletion in sorted marker-negative cells.

As DIV-MARIS robustly isolates neurogenic cell populations of interest, we extracted RNA from sorted populations at four developmental time points for whole-transcriptome sequencing. Principal component analysis demonstrates that variance between samples is primarily due to developmental time and cell type of origin (Fig. S4). The resulting cell-type specific gene expression atlas quantitatively assesses neurogenic transcription in five distinct neurogenic cell populations (enriched and depleted) across 4 developmental time points covering major neurogenic events (Fig 1A, B).

### Cell type-specific expression of protein-coding genes during neurogenesis

In addition to purity, we evaluated sorting *specificity* by assessing gene expression of the five cell type marker genes (*ind, vnd, pros, elav,* and *repo*) across the sorted populations in terms of normalized counts (File S1). In all cases, strong enrichment of marker gene expression levels in the marker-enriched compared to the depleted samples was observed as expected (Fig. 2A, S5). For example, the high and near-exclusive enrichment of *repo* transcript in purified glia demonstrates sorting effectiveness of DIV-MARIS when using a highly specific and exclusive marker (Fig. 2A, S5E). Similarly, *elav* transcript levels are highly enriched in purified neurons compared to glia (Fig. 2A, S5D), while lower levels can be detected in early neuroblasts and columnar material, which is in line with observations that the common neuronal marker *elav* is transiently expressed pan-neurogenically at the onset of differentiation (Berger et al. 2007). The columnar markers *vnd* and *ind* mark distinct columnar neurogenic territories that each give rise to neuroblasts, neurons and glia. Accordingly, while *vnd* and *ind* transcripts are largely exclusive to their respective neurogenic columns, each is detectable to some degree in neuroblasts, most likely because early neuroblasts stem from one of the respective neurogenic columns co-purified by FACS (Fig 2A, S5A-C). Interestingly, normalized counts for *vnd* are higher than *ind,* which likely reflects that the ventral column generates more neuroblasts in the first waves of delamination compared with the intermediate column (Doe 1992).

**Fig. 2.**
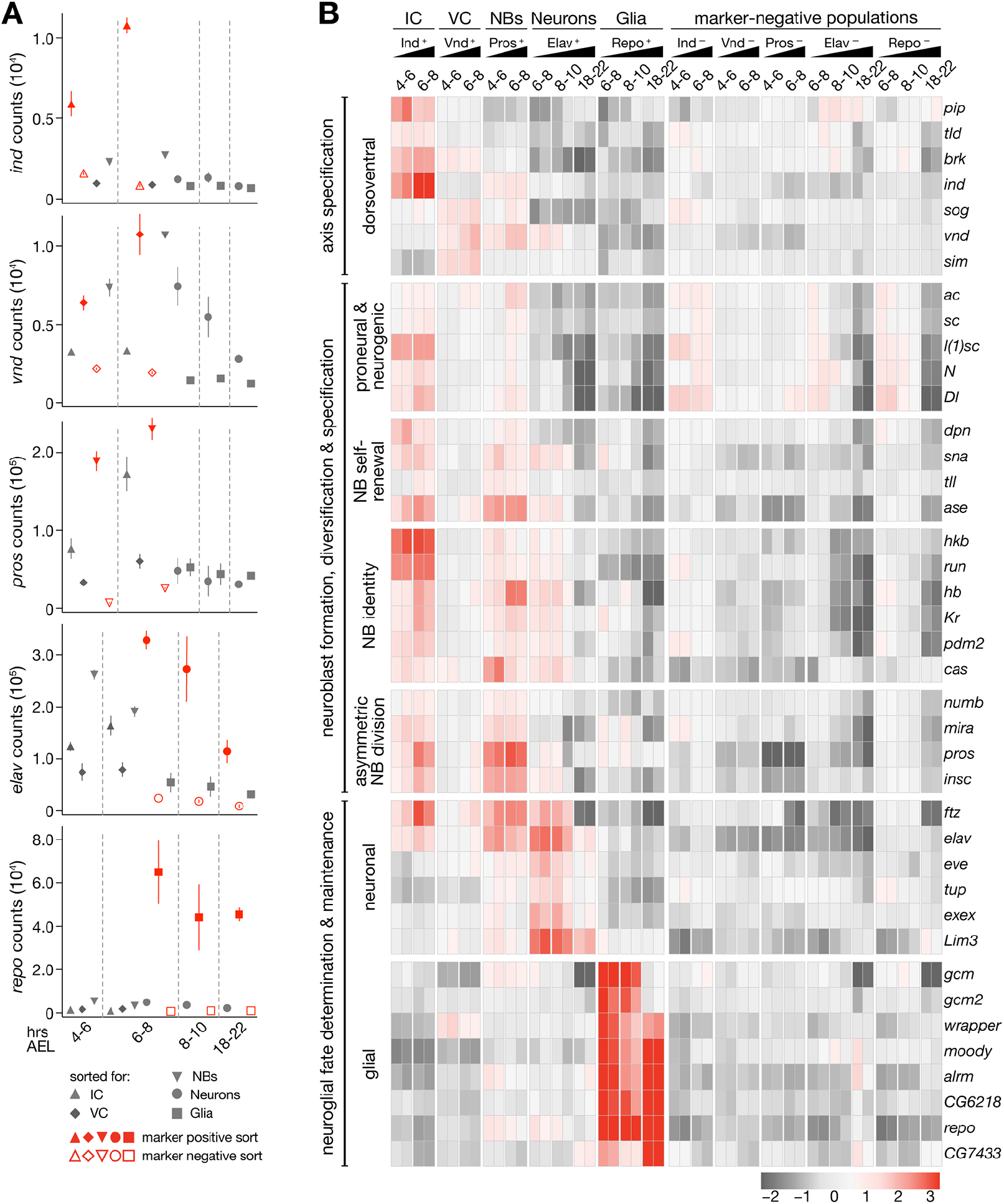
Defining mRNA signatures of neuroglial cell types. (A) Normalized expression values for each marker gene used for FACS (*ind, vnd, pros, elav,* and *repo*) across sorted samples. (B) Heat map of expression profiles of Drosophila nervous system genes. Row mean-centred expression values calculated via variance-stabilizing transformation of gene-level RNA-seq counts (scale = log_2_ ratio of row mean).

To validate cell type-specific gene expression, we examined genes with known neurogenic roles (Ohlen & Doe 2000a; Skeath & Thor 2003; Sousa-Nunes et al. 2010; Crews 2010; Sandler & Stathopoulos 2016) and confirmed specificity of mRNA expression in cell types previously associated with gene function (Fig. 2B). *Exex,* for example is a homeodomain transcription factor required in motor neurons that are projecting to ventral somatic muscles (Santiago et al. 2014) and we find it exclusively in young neurons (Fig. 2B). While markers of neuroblast identity were not only enriched in neuroblasts, but depleted in the differentiated cell types neuroblasts give rise to (neuronal and especially glial), neurogenic column marker expression was often maintained in neuroblasts, highlighting that neuroblasts retain columnar identity after delamination, as they are adopting column-specific fates (Doe 1992).

To systematically uncover protein-coding genes that demarcate columnar and cell type identities in nervous system development, we queried for genes expressed similarly to known neurogenic genes by Pearson correlation (*r* > 0.9). We uncovered 753 additional genes (summarized in File S2) and though many have no known association with embryonic neurogenesis, *in situ* screens annotating expression using controlled anatomical imaging vocabulary (ImaGO, Hammonds et al. 2013; Tomancak et al. 2002; Tomancak et al. 2007) indicate that this gene set is indeed specifically expressed in components of the developing nervous system. For example, the most enriched ImaGO terms for this gene set include “ventral nerve cord primordium”, “brain primordium”, and “ventral nerve cord” (Fig. S6A). GO analysis reveals the most enriched molecular function for this gene set appears to be “DNA binding”, and the most enriched biological processes are “chromosome organization” and “nucleic acid metabolic process” (Fig. S6C-D). Furthermore, protein domains enriched among the proteins specifically expressed in compartments of the developing nervous system are enriched for histone folds, chromatin interaction domains and sequence specific DNA interaction domains, such as zinc fingers and homeobox domains (Fig S6D).

We were surprised that one quarter of the genes deployed similarly to known neurogenic marker genes remain largely unstudied (199 ‘computed genes’) and though many of these candidates lack *any* described function, more than 62% can be directly mapped to human homologs and some have even been linked to nervous system function.

We focused on a subset (40) of these genes, which were predicted to be expressed in neuroglial cell types with clear spatiotemporal specificity (Fig S7A). In concordance with DIV-MARIS predictions, RNA *in situ* hybridization data (Hammonds et al. 2013; Tomancak et al. 2002; Tomancak et al. 2007) confirms that a selection of these candidate genes mark specific subsets of cells in the developing nervous system (Fig. S7B).

Thus, DIV-MARIS reliably captures and uncovers cell type-specific gene expression dynamics during embryogenesis. As many of the specifically expressed genes encode known and predicted transcription factors and signaling pathway components (File S2), this cell type-specific expression map identifies new regulatory nodes that likely play central roles in the specification and differentiation of neuroglial cell types.

### Specific expression and properties of long noncoding RNAs along the neuroglial lineage

To explore lncRNA expression during early neurogenesis, we first identified nervous system-specific lncRNAs by calculating enrichment of expression in marker-positive versus marker-depleted samples at each time point using DESeq2 (Love et al. 2014; log_2_FC >1.0, *padj* <0.05). We found 325 such lncRNA candidates (File S3) and evaluated them according to several criteria, including spatiotemporal regulation through neurogenesis, expression above an abundance threshold in at least one cell type (TPM >300), absence of sense overlap with a protein-coding gene, and transcript boundaries consistent with lncRNA annotations. Applying these stringent criteria, we selected 13 high-confidence lncRNA candidates that are strongly and specifically expressed in a variety of cell types of the *Drosophila* nervous system (Fig. 3, S8).

**Fig. 3.**
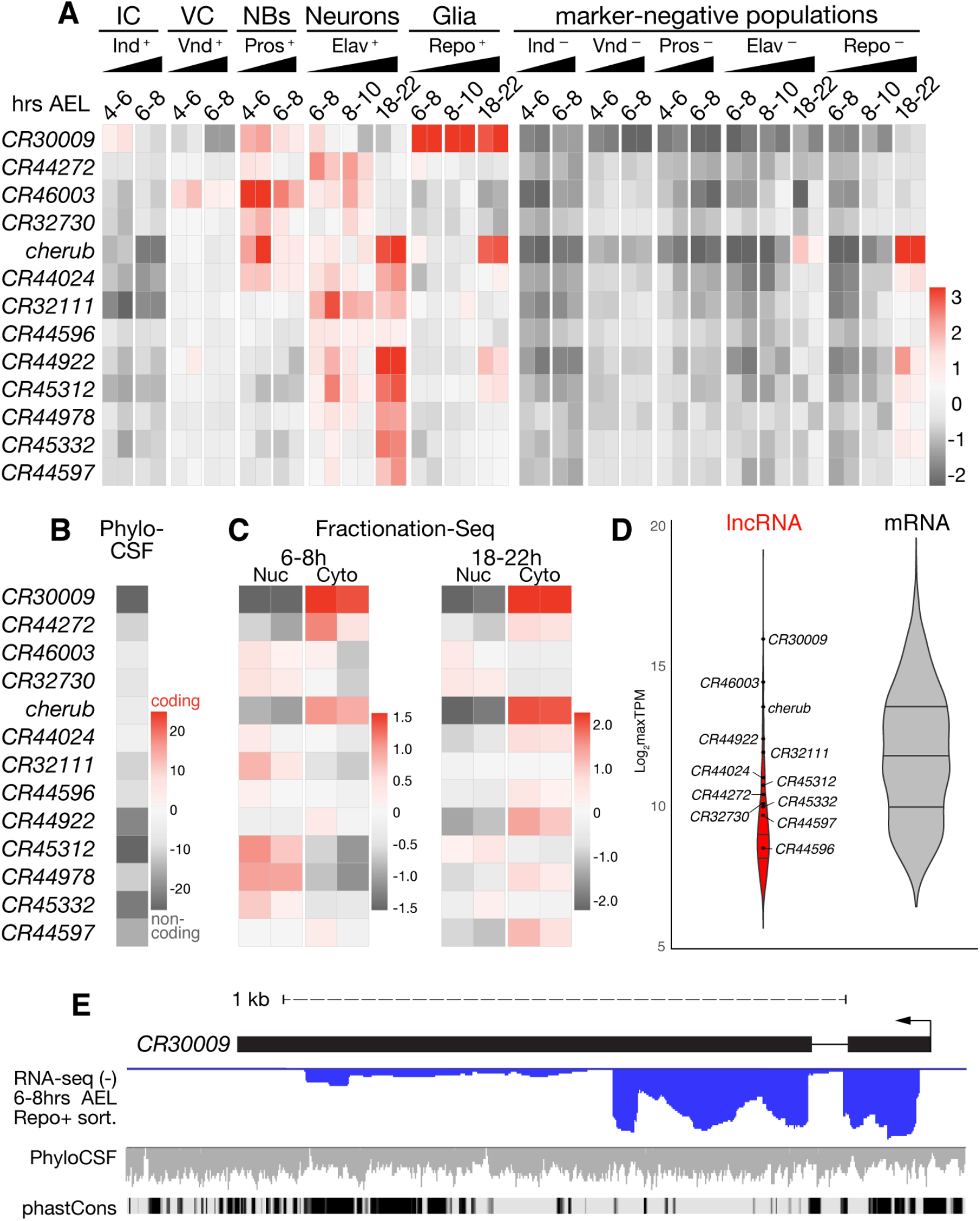
Neuroglial lncRNAs are highly regulated transcripts. (A) Row mean-centered expression values of lncRNAs in marker-enriched and -depleted samples calculated via variance-stabilizing transformation of gene-level RNA-seq counts (scale = log_2_ ratio of row mean). (B) PhyloCSF scores (ScorePerCodon) for the putative ORF with highest coding potential within each transcript. Scale is roughly one standard deviation above the mean score of coding regions (very high coding potential) down to one standard deviation below the mean of non-coding regions (very low coding potential). (C) Row mean-centered expression profiles in 6-8h and 18-22h embryo nuclear and cytoplasmic fractions generated by Fractionation-Seq; values as in (A). (D) Violin plot showing distribution of maximum TPM (maxTPM) values for all lncRNAs (red; n = 325) and mRNAs (grey; n = 3835) differentially expressed (log_2_FC >1.0, *padj* <0.05) between any marker-positive and marker-negative cell type; lncRNAs presented in (A) are highlighted. (E) *CR30009* genomic locus showing stranded RNA-seq data from sorted glia at 6-8hrs AEL (negative strand; blue), overlay of smoothed PhyloCSF scores of individual codons in each of three frames (horizontal line is 0), and conservation among Drosophilids (phastCons).

To assess spatiotemporal expression of these lncRNAs, we calculated their relative abundance among all cell types and collection bins. The lncRNAs are depleted in marker-negative non-neurogenic cells and exhibit dynamic spatiotemporal enrichment in specific marker-positive cell types (Fig. 3A). Strikingly, while we found very few lncRNAs with distinct expression in the earlier and more naïve intermediate and ventral columns, specific lncRNA deployment can be readily observed in more mature and differentiated cell types, such as neuroblasts, neurons, and/or glia, indicating that lncRNA expression is a hallmark of differentiated cells more so than of primordia.

To confirm that these transcripts are *bona fide* lncRNAs, we evaluated the coding potential of each via Phylogenetic Codon Substitution Frequency (PhyloCSF) (Lin et al. 2011). Each lncRNA locus exhibits a total PhyloCSF score below zero across all frames, consistent with a complete lack of coding potential (Fig. 3B). Given that some lncRNAs have been shown to exhibit variable subcellular localization with localized functions (L.-L. Chen 2016), we assessed the general subcellular expression of these transcripts via Fractionation-Seq. Briefly, we generated a subcellular reference transcriptome of the cytoplasmic and nuclear compartments of 6-8h and 18-22h embryos and examined abundance of each of these lncRNA transcripts between these fractions. Intriguingly, the 13 lncRNAs exhibit distinct subcellular localization patterns with varying degrees of nuclear/cytoplasmic restriction (Fig. 3C), ranging from almost exclusive cytoplasmic (e.g. *CR30009* & *cherub*) to almost exclusive nuclear detection (e.g. *CR45312*), including instances where location appears temporally regulated (e.g. *CR44978*).

To assess lncRNA abundance relative to other transcripts (noncoding *and* protein-coding) in the neurogenic cell types, we normalized read counts for each transcript in each sample (TPM, File S4). The maximum expression score for lncRNAs across cell types (maxTPM) shows that while expression varies among lncRNAs, they are generally not *lowly* expressed; rather, lncRNA expression is well within the range of what may be expected for protein-coding genes significantly regulated during neurogenesis (Fig 3D). That these lncRNAs are *bona fide* regulated transcripts is further supported by specific splicing, which is observed for several of the neurogenic lncRNAs (Figs 3E, S8). Thus, these lncRNAs are unlikely to be merely byproducts of spurious transcription, rather they are subject to regulated expression, RNA processing, and controlled export, which supports a potential role in neurogenesis.

One intriguing example of a lncRNA demonstrating specific expression over the course of early neurogenesis is *CR30009.* This lncRNA shows increased expression in the early intermediate column and in neuroblasts, but is most highly enriched in glial cells during all assayed time windows (Fig. 3A). Furthermore, *CR30009* is spliced and primarily exported to the cytoplasm (Figs 3C,E), features indicative of specific co-and post-transcriptional regulation. However, *CR30009* has the lowest coding potential out of all tested lncRNAs – its PhyloCSF score per codon (−42.647) is more than three standard deviations below the mean for noncoding regions in *Drosophila* (−18.7 ± 7.2, Fig. 3B). Furthermore, *CR30009* is one of the most highly abundant transcripts in glia – noncoding or protein-coding (log_2_ maxTPM = 15.75, Fig. 3D, File S3) – which underscores the *potential* functional importance of *CR30009* in gliogenesis. Notably, this lncRNA appears to exist predominantly as an unannotated short isoform and exhibits regions of high non-coding sequence conservation among *Drosophilids* within the first exon and at the 3’ end of the transcript (Fig. 3E).

A second example, *CR43283* (also known as *cherub*), exhibits dynamic temporal regulation. Expression of *cherub* is strongly enriched in the earliest neuroblasts at 4-6h, but enrichment quickly wanes in later neuroblasts (6-8h); however, over time *cherub* becomes specifically expressed being strongly enriched in differentiated neurons and glia by the end of neurogenesis at 18-22h AEL (Fig. 3A). We note that enriched expression of the lncRNA in Elav- and Repo-negative samples may be contributed by cherub-positive glia in the neuron–depleted fraction and cherub-positive neuroblasts/neurons in the glia-depleted fraction. *cherub* is also specifically localized to the cytoplasm throughout embryogenesis, is clearly spliced, but harbors no coding potential (Figs. 3B-C, S8D).

*CR32730* is first detected in 4-6h neuroblasts and is moderately enriched at 8-10h in the neuronal, but not in the glial population (Fig. 3A). *CR32730* is transcribed antisense to the intron of *CG9650* (Fig S8C), a putative neurogenic transcription factor that has been implicated in CNS development (McGovern et al. 2003). However, *CR32730* appears to be transcribed independently of *CG9650,* which is lowly expressed in early neuroblasts according to DIV-MARIS (Fig. S8C), suggesting that their roles could be independent. Fractionation-Seq predicts that *CR32730* is moderately enriched in the nuclear fraction in early and late embryos (Fig. 3C).

Expression of another lncRNA, *CR46003,* is first detected in the ventral column and is most highly enriched in early neuroblasts, but expression persists in neuroblasts and early neurons (Fig. 3A-C). *CR46003* is among the most abundant lncRNAs in our dataset and does not exhibit clear subcellular enrichment in either early or late embryos (Fig 3 C-D). Intriguingly, the transcription start site of *CR46003* is antisense to *CR46004,* which contains a miRNA implicated in behavior (Picao-Osorio et al. 2017) (Fig. S8B).

*CR44024* expression is first enriched in early neuroblasts and persists through neuronal differentiation, and is predicted to be excluded from the intermediate and ventral columns and glia (Fig 3A). This lncRNA is not predicted to exhibit distinct subcellular localization in early (6-8h) embryos, but is moderately enriched in the cytoplasm at the end of embryogenesis (18-22h, Fig 3C). *CR44024* is also one of the highly expressed lncRNAs in our dataset and is on par with protein-coding genes (Fig 3D). The transcript is intergenic, and appears to be spliced, although not in accordance with its annotated transcript model (Fig. S8E).

In summary, DIV-MARIS predicted spatiotemporal expression of a number of lncRNAs during neurogenesis. Through the application of stringent criteria, we refined this list to a high-confidence selection of noncoding transcripts with diverse predicted expression patterns and properties. To confirm these predictions for several lncRNA candidates, we first visualized their expression in the context of a whole developing embryo.

### Neurogenic lncRNAs mark specific neuroglial subsets of cells

To visualize lncRNA expression, we performed multiplex RNA-FISH (Kosman et al. 2004) against the five examples discussed above (*CR30009, cherub, CR46003, CR32730,* and *CR44024*) together with neurogenic marker genes. Remarkably, RNA-FISH reveals exquisite spatiotemporal specificity of lncRNA expression for each of the lncRNAs tested.

*CR30009* – predicted by DIV-MARIS to be highly enriched in glia – is indeed co-expressed with *repo* as expected in clusters of glial cells as early as stage 9/10 (Figs 4A-B, S10). *CR30009* remains co-expressed with most repo-expressing cells through stage 13/14 (Fig. 4C-D). However, timing of *CR30009* expression suggests it to be independent of *repo,* indicating that the lncRNA constitutes an earlier marker of the glial lineage than currently known. While most repo-positive cells also express *CR30009* in stage 9-12 embryos, the lncRNA is largely expressed in small puncta within other cells in the ventral nerve cord and brain that are likely to be neuroblasts (Figs 4A-B, S10). Accordingly, DIV-MARIS predicts *CR30009* expression in 4-6h and 6-8h pros-positive cells (Fig. 3A, S11). It is feasible, therefore, that the lncRNA *CR30009* constitutes the earliest neuroblast marker of the glial lineage to date, which accumulates into larger, brighter foci during early phases of glial differentiation (Fig. S10, S11).

**Fig. 4.**
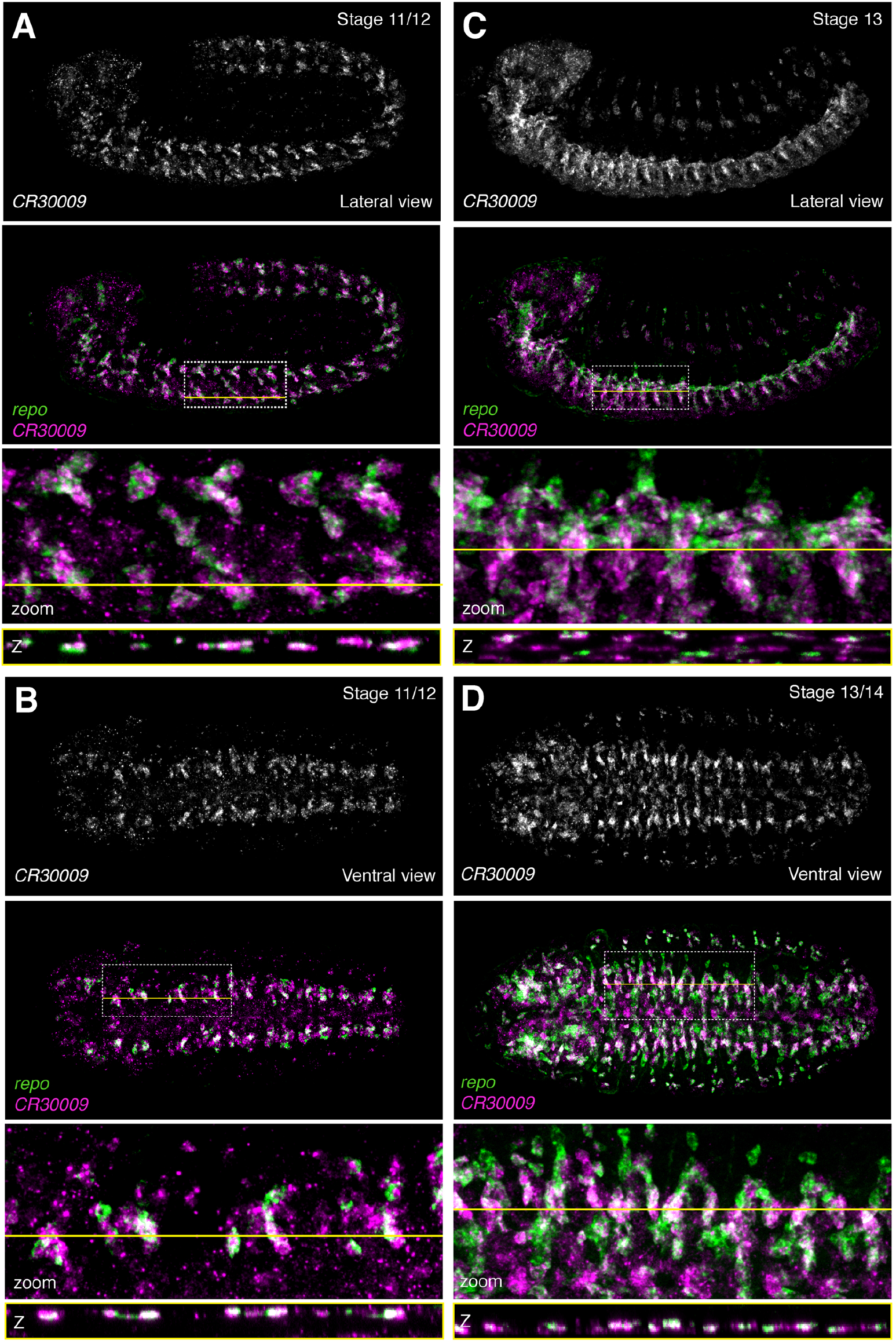
The lncRNA *CR30009* is expressed in glial subsets. RNA fluorescent *in situ* hybridization (RNA-FISH) against *CR30009* and the glial marker *repo.* (A) Lateral view, stage 11/12. (B) Ventral view, stage 11/12. (C) Lateral view, stage 13. (D) Ventral view; stage 13/14. Top, *CR30009* alone; below, merge of *CR30009* (magenta) with *repo* (green). Dashed white box indicates region of interest (ROI) and yellow line indicates Z-slice through ROI. Second from bottom: zoom in of ROI. Bottom: Slice through Z-stack as indicated by yellow line.

RNA-FISH against the lncRNA *cherub* revealed strong spatiotemporal regulation of *cherub* broadly in accordance with DIV-MARIS, which predicted *cherub* to be strongly and specifically enriched in early (4-6h) neuroblasts, and late (18-22h) neurons and glia (Fig. 3A).

We observed clear induction of *cherub* expression within six small clusters of cells in the ventral nerve cord during stage 12, each of which also expresses *pros* (Fig 5A) and to a lesser degree, *elav* (Fig 5B), both of which is in line with *cherub* constituting a neuroblast marker. During stage 13, *cherub* is seen in several additional clusters in the brain (Figs 5B, S12A-B). By stage 14-15, *cherub* is very strongly expressed in multiple defined *pros* neuroblast clusters, but might be excluded from mature neurons and glia (Figs 5C-D, S12C-D, S13), and remains strongly expressed through the remainder of embryogenesis (stage 16-17, Fig S13B), in line with DIV-MARIS predictions (Fig 3A).

**Fig. 5.**
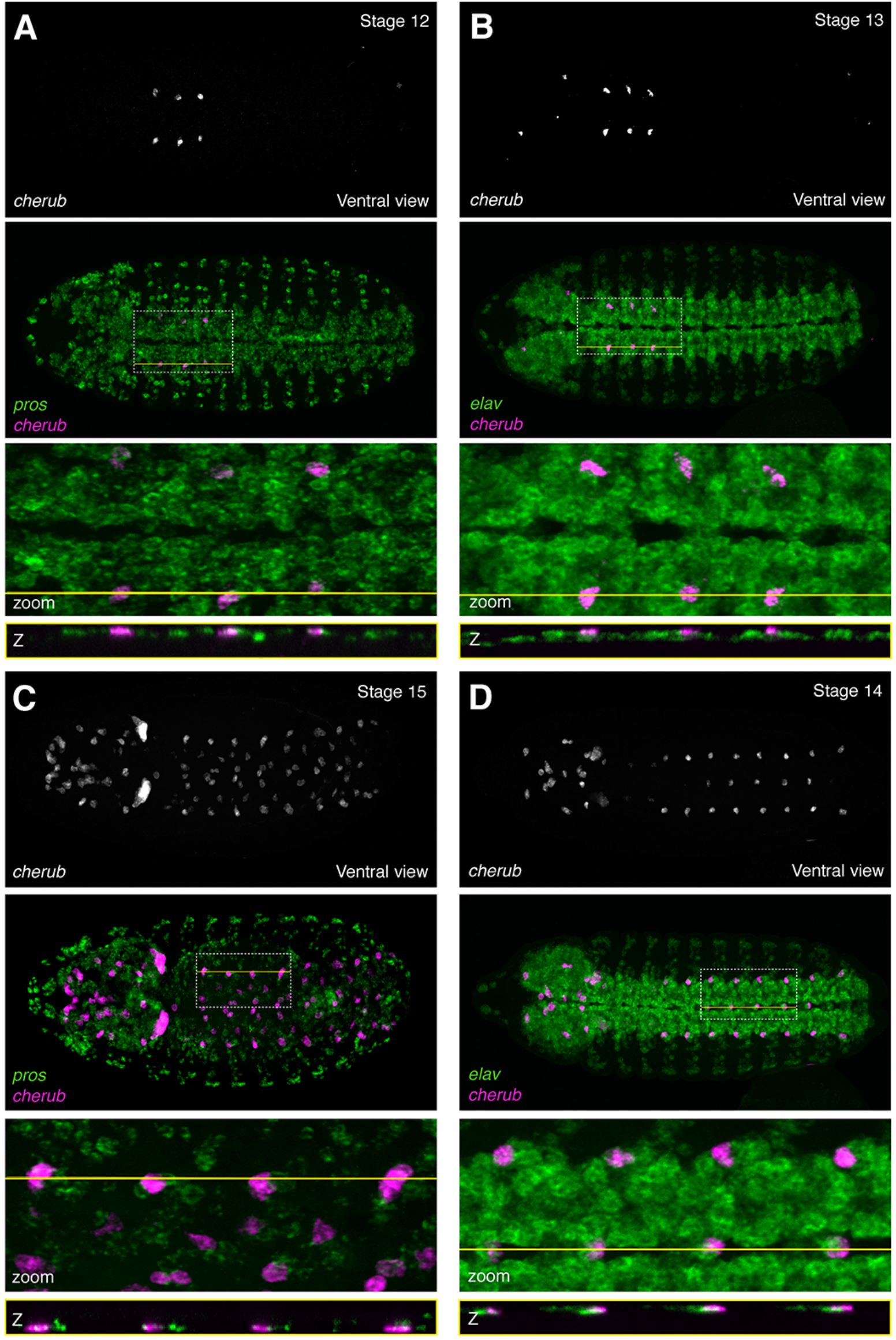
The lncRNA *cherub* is expressed with strict spatiotemporal specificity primarily in a subset of neuroblasts. RNA fluorescent in situ hybridization (RNA-FISH) against *cherub,* the neuroblast marker *pros,* and the neuronal marker *elav.* Ventral view. (A) *cherub* with *pros*; stage 12. (B) *cherub* with *elav;* stage 13. (C) *cherub* with *pros;* stage 15. (D) *cherub* with *elav;* stage 14. Top: *cherub* alone. Second from top: *cherub* (magenta) overlaid with marker (green). Dashed white box indicates region of interest (ROI) and yellow line indicates Z-slice through ROI. Second from bottom: zoom in of ROI. Bottom: Slice through Z-stack as indicated by yellow line.

DIV-MARIS predicts similar spatiotemporal expression of *CR46003* and *CR32730* in neuroblasts and neurons (Fig. 3A). Indeed, RNA-FISH reveals very similar patterns of expression of the two lncRNAs. *CR46003* is the earliest expressed among all lncRNAs tested here and is detected in a small cell cluster already at stage 5-6 (Fig S14). By stage 9-10, punctate expression of *CR46003* appears in defined *pros*-expressing clusters along the embryonic ventral midline (Figs 6A), in agreement with the DIV-MARIS-predicted enrichment in cells of the ventral column and neuroblasts at 4-6 and 6-8hrs AEL (Fig. 3A). *CR46003* expression expands to a greater number of cells within and beyond the ventral nerve cord and brain from stage 11-13, many of which also express *pros* (Fig 6B, S15B-C) and some express *elav* as well. (Fig.S16). As predicted by DIV-MARIS, RNA-FISH demonstrated that *CR32730* follows a very similar pattern of expression to *CR46003* from stage 9-10 (Figs 6D, S15D) through stage 13 (Figs 6, S15E-F).

**Fig. 6.**
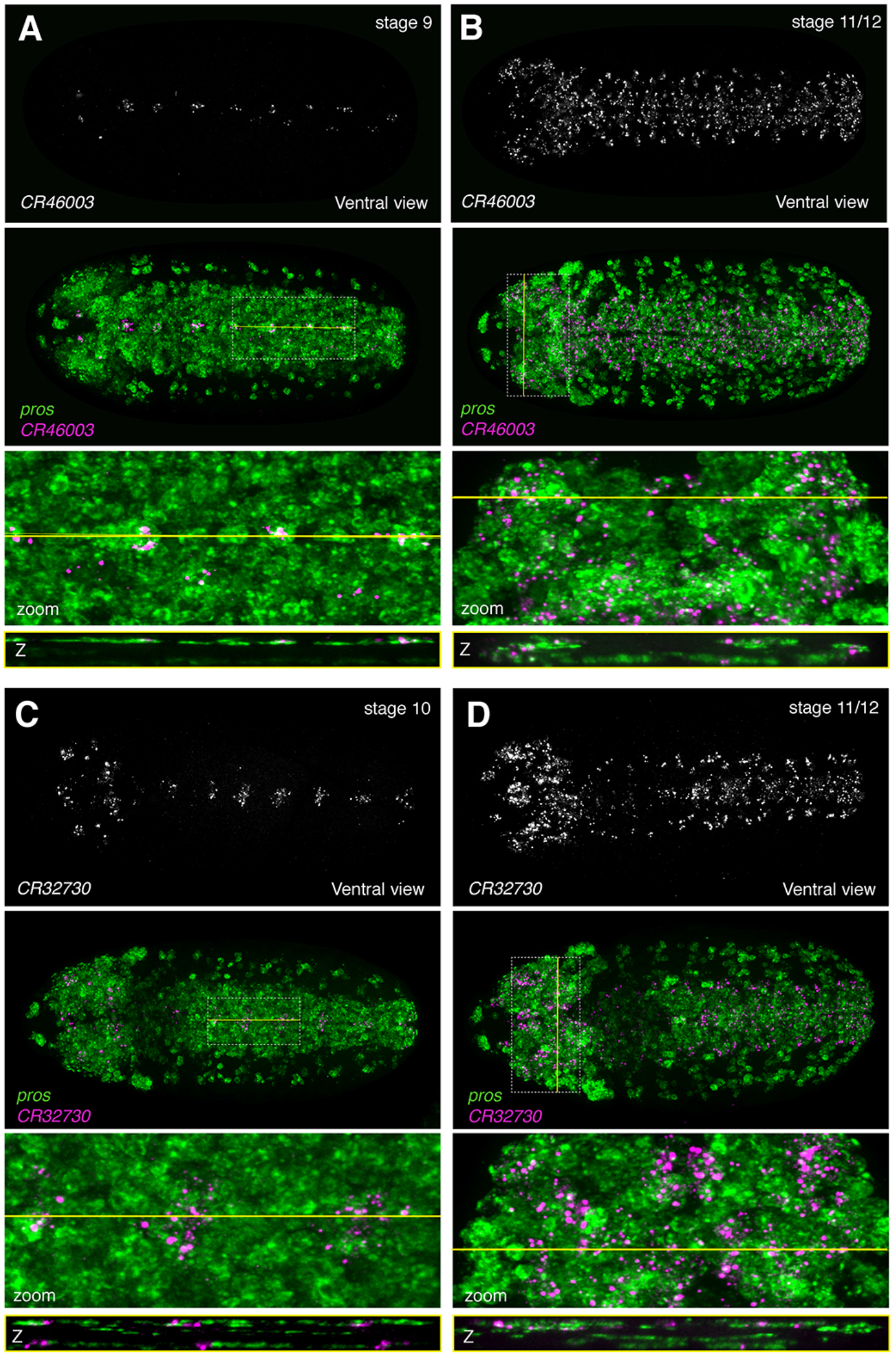
The lncRNAs *CR46003* and *CR32730* are expressed with similar spatiotemporal specificity in a subset of neuroblasts. RNA fluorescent in situ hybridization (RNA-FISH) against *CR46003* and *CR32730* together with the neuroblast marker *pros.* Ventral view. (A) *CR46003*; stage 9. (B) *CR46003;* stage 11/12. (C) *CR32730*; stage 10. (D) *CR32730;* stage 11/12. Top: lncRNA alone. Second from top: lncRNA (magenta) overlaid with *pros* (green). Dashed white box indicates region of interest (ROI) and yellow line indicates Z-slice through ROI. Second from bottom: zoom in of ROI. Bottom: Slice through Z-stack as indicated by yellow line.

While we were not able to detect the transient *CR44024* expression in early (stage 9-10) prospero positive neuroblasts as predicted by DIV-MARIS, we did however observe that the lncRNA exhibits highly dynamic temporal regulation. At stage 12, *CR44024* is induced within small *elav-positive* clusters flanking the midline (Fig. S17). Starting at stage 13, *CR44024* is expressed more much broadly, yet still restricted to subsets of *elav-* and pros-expressing cells within the ventral nerve cord and central brain (Fig. 7).

**Fig. 7.**
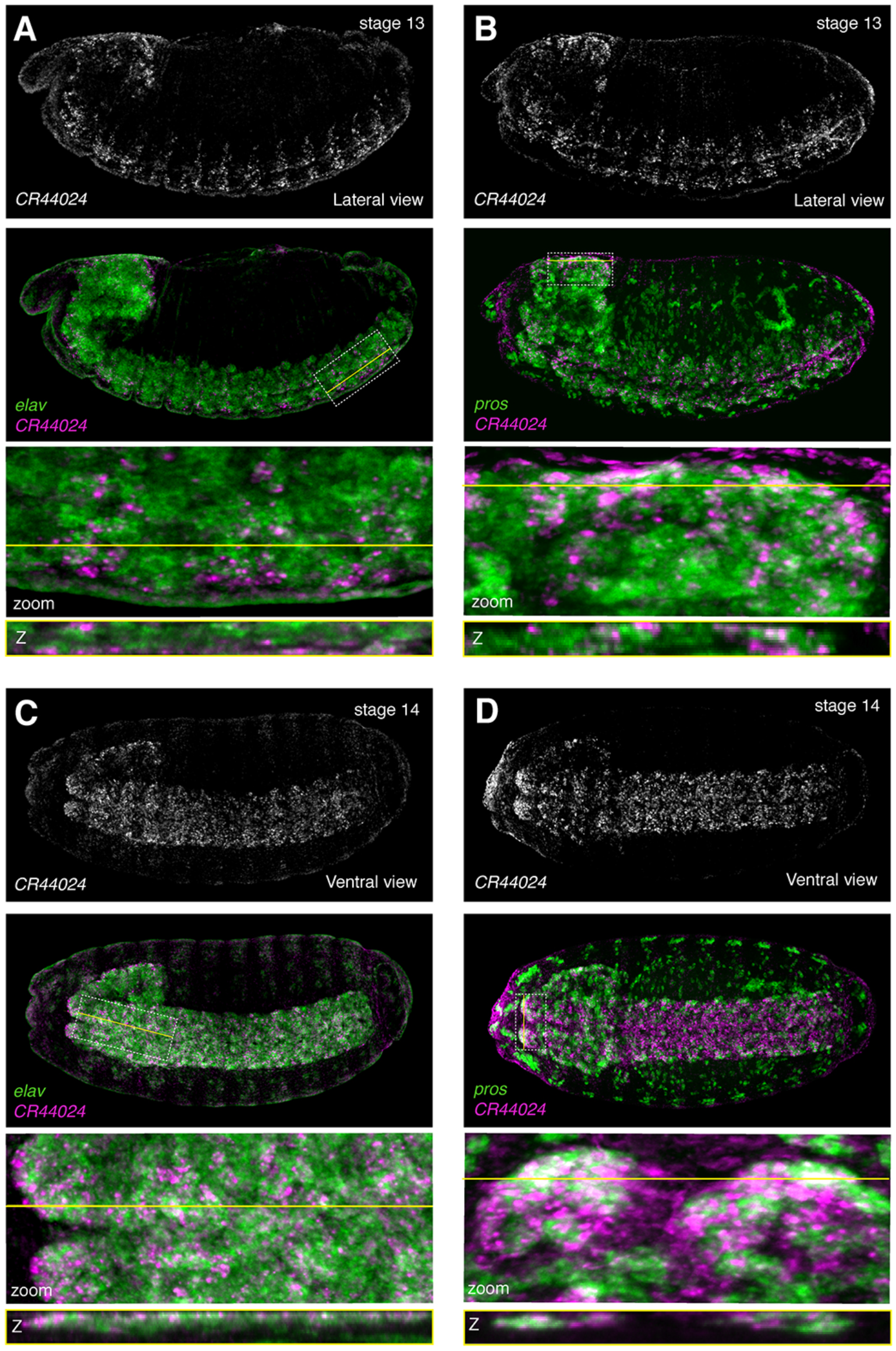
The lncRNA *CR44024* is expressed later in embryogenesis in neuronal subsets. RNA fluorescent in situ hybridization (RNA-FISH) against *CR44024,* the neuroblast marker *pros,* and the neuronal marker *elav.* (A) *CR44024* and *elav;* lateral view; stage 13. (B). *CR44024* and *pros;* lateral view; stage 13. (C). *CR44024* and *elav;* ventral view; stage 14. (D). *CR44024* and *pros;* ventral view; stage 14. Top: *CR44024* alone. Second from top: *CR44024* (magenta) overlaid with marker (green). Dashed white box indicates region of interest (ROI) and yellow line indicates Z-slice through ROI. Second from bottom: zoom in of ROI. Bottom: Slice through Z-stack as indicated by yellow line.

Lastly, we assessed the subcellular localization of individual lncRNAs. For example, Fractionation-Seq (Fig. 3C) predicted *CR30009* and *cherub* to be predominately cytoplasmic. This is supported for both lncRNAs by high-resolution confocal microscopy, as both transcripts are primarily detected in the cytoplasm (Fig S18A-B). *CR46003* and *CR32730* both showed a slight bias for nuclear localization by Fractionation-Seq, which was confirmed by microscopy as both lncRNAs clearly stain within the nucleus, though it should be noted that subnuclear puncta are observed (Fig. S18C-D). Similarly, *CR44024* appears to be restricted primarily to the nucleus in the ventral nerve cord at stage 14 (Fig. S18E), matching the prediction.

The identification of such complex, yet specific expression patterns highlights the importance of tissue- and cell type-specific expression analysis. Whole embryo studies, for example, not only lack spatial resolution, but expression signatures – even of highly expressed genes – may be lost if their expression is specific to a small-enough subset of cells. Here, we provide a map for the cell type-specific expression of coding, as well as noncoding RNAs over the course of embryonic neurogenesis in the developing *Drosophila* embryo. While hundreds of coding and dozens of lncRNAs are deployed with specific spatial and temporal dynamics, it should be noted that direct imaging of expression in spatial context can reveal nuances of expression that is beyond the resolution of many cell type-specific genomic approaches.

## Discussion

Complex tissues are defined by the intricate interplay of individual cell types that differ in their gene expression programs. Tissue culture has long been an important tool for the genome-wide investigation of cellular responses as it avoids much of the heterogeneity inherent to living tissues. Unfortunately, it is often precisely this heterogeneity and the dynamic contacts between cells and tissues that often shape cellular identities and transcriptomic responses. Hence, to determine the gene regulatory programs that drive complex organismal development, it is crucial to (i) preserve the cellular interactions *in vivo,* (ii) acquire genome-wide transcriptomic data with spatial and/or cell type resolution, and (iii) assure temporal resolution.

### DIV-MARIS to investigate global cell type-specific gene expression dynamics

To investigate the transcriptome dynamics over the course of neurogenesis from primordial to neuronal and glial identities, we developed a method to isolate specific cell types from *Drosophila* embryos with resolution in time and space. DIV-MARIS is widely applicable and can be employed for spatiotemporal transcriptional profiling of basically any cell type of interest in the *Drosophila* embryo and other complex tissues, as long as markers allowing for sorting a cell type of interest are available (i.e. appropriate antibodies or transgenic markers e.g. *enhancer-reporter* constructs). DIV-MARIS employs chemically cross-linking of the cellular material, thus assuring that the developmental *status-quo* is preserved and elaborate sorting strategies based on multiple markers could be devised to fine-tune the sub-population selection one wishes to purify (Molyneaux et al. 2015).

Here, we purified fixed cells based on markers of specific neurogenic cell populations in the early *Drosophila* embryo. DIV-MARIS faithfully resolved known expression patterns of neurogenic protein-coding genes, but also identified cell type specific expression of additional genes with yet unknown neurogenic functions – while neuroglial expression was confirmed by *in situ* hybridization for a few dozen mRNAs, hundreds more are predicted to exhibit spatiotemporal expression over the course of early neurogenesis. This compendium lays the groundwork for a comprehensive understanding of the mechanisms driving early neurogenesis and given that many of the spatiotemporally expressed genes encode regulatory factors such as transcription factors and signaling molecules, careful examination of their neurogenic roles will be required.

### Identification of spatiotemporal lncRNA expression

This study has identified many cell type-specific lncRNAs with potential neurogenic function. We emphasize that this is not yet an exhaustive list of lncRNAs expressed in the nervous system, as our filtering criteria were conservative. Instead, we focused on a high-confidence set of 13 lncRNAs with a variety of expression and transcript characteristics. Given that these noncoding transcripts are (i) temporally expressed in specialized cell types and subtypes of the nervous system, (ii) moderately-to-highly abundant and (iii) often exhibit hallmarks of RNA processing (such as splicing and nuclear-cytoplasmic shuttling), these lncRNAs appear to be subject to regulated expression rather than being by-products of spurious transcription.

Notably, we did not identify any lncRNAs with expression *restricted* to the early neuroectodermal columns. According to DIV-MARIS, there is some enrichment of *CR30009* in the intermediate column and *CR46003* in the ventral column (Fig. 3A), but since the respective territorial markers of the ventral and intermediate columns are still detectable in the neuroblast progenitors, this expression may be specific to neuroblasts, in which higher enrichment is observed. High and specific lncRNA expression appears to be a feature of differentiating and differentiated cell types of the nervous system, rather than of primordial territories.

Multiplex RNA-FISH shows that lncRNAs often exhibit a high degree of cell type specificity. Though co-expression is generally detected with cell type-specific markers as predicted by DIV-MARIS, we could observe much more nuanced spatiotemporal lncRNA regulation than we could have predicted – the noncoding transcripts investigated here tended to be expressed in highly specific subsets of neurogenic cell types (Figs 4–7). It is therefore feasible that these lncRNAs perform highly specialized functions in subsets of cells contributing to discrete regions of the nervous system.

For example, *CR46003* and *CR32730* are the first lncRNAs that appear to specifically mark midline and midline-proximal structures (Fig. 6, S15). Given the midline’s highly specialized role as a signaling and organizing center (Wheeler et al. 2006; Crews 2010; Zhou et al. 1995), it is intriguing to speculate that such lncRNAs may help shape the midline fates. While lncRNAs were enriched in a variety of neurogenic populations, *CR30009* was consistently and highly enriched in repo-positive glia and to some degree in pros-expressing neuroblasts (Figs 4, S10-11). It is feasible that *CR30009* may play a role in the priming of glial fates from the earliest stages of differentiation, possibly mediating the transition from neuroblasts and GMCs to specifically the glial fate. As most glia in the *Drosophila* embryonic CNS originate from the lateral column, it will be of interest if *CR30009* expression and function may be limited to glia of the lateral neurogenic ectoderm, or if it is present in ventral column-derived glia as well.

Are these lncRNAs functional? *cherub* serves as a nice example arguing that several of them likely are. The lncRNA *cherub* was recently identified as a highly up-regulated transcript in neuroblast-derived tumors in larvae (Landskron et al. 2018). In larvae, *cherub* is asymmetrically inherited by the self-renewing neuroblast to allow fate progression of the sibling cell and *cherub*’s specific predicted enrichment in embryonic neuroblasts (Fig. 3A) indicates that this lncRNA could exhibit a similar function in the early embryo. However, the precise temporal regulation of *cherub* was surprising, as RNA-FISH identified its presence not in early, but in differentiating and fully differentiated neurons and glia by the end of embryogenesis (Figs 5, S12-13).

Intricate spatio-temporal expression regulation is a hallmark of many lncRNAs and not only in the *Drosophila* embryo as well as generally (Wilk et al. 2016; Karaiskos et al. 2017; Landskron et al. 2018). Various lncRNAs have been demonstrated to play diverse biological roles – nuclear and cytoplasmic – from integral parts of riboprotein complexes, to regulating dosage compensation, to affecting genome topology. LncRNA complexity was reported to be especially pronounced in the nervous system and even early stages of embryonic neurogliogenesis appear to be no exception. However, the challenge clearly remains to unravel the neurogenic roles of these putative noncoding regulators, and the molecular mechanisms by which they act. This study represents a valuable resource for understanding transcriptome complexity in the emerging nervous system and it lays the basis for further studies into the mechanisms by which non-coding genes, but also hundreds of specifically deployed coding genes shape nervous system development.

## Materials and Methods

### Fly lines

See Supplementary Materials and Methods.

### FACS purification and RNA isolation using DIV-MARIS

Briefly, embryos were dissociated into single-cell suspensions, cells were fixed in 4% formaldehyde. Fixed cell suspensions were immunostained under RNase-free conditions and FACS-purified using a FACS-AriaII cell sorter (BD Biosciences). Marker-enriched and -depleted cell populations were collected in biological duplicates. FACS-purified cells were subject to cross-link reversal and proteinase K digestion prior to RNA isolation. Additional experimental details for DIV-MARIS are provided in the Supplementary Materials and Methods; primary and secondary antibodies used in this study are listed in Table S1.

### Nuclear-cytoplasmic fractionation

Cytoplasmic and nuclear extracts were isolated from whole *Drosophila* embryos via detergent-based hypotonic lysis for RNA isolation. Additional experimental details are provided in the Supplementary Materials and Methods.

### Quantitative RT-PCR (qPCR)

qPCR was performed using standard SYBR Green, using the BioRad CFX96 Touch™ Real-Time PCR Detection System. Additional information available in the Supplementary Materials and Methods; qPCR primer sequences are listed in Table S2.

### Library preparation and RNA-sequencing

All RNA-seq libraries were constructed using the NuGEN Ovation *Drosophila* RNA-Seq System with 10 ng – 100 ng total RNA input. Library concentration was quantified using the Qubit™ dsDNA HS Assay (Thermo, Q32854) and quality was determined on a BioAnalyzer™ using Agilent High Sensitivity DNA Kits (Agilent, 5067-4626). All libraries were sequenced on the Illumina HiSeq4000 at a mean depth of 62.5 million 75bp paired-end reads per sample. RNA-seq datasets generated for this study are detailed in Tables S5 and S6. All RNA-seq data has been deposited into the NCBI Gene Expression Omnibus (GEO) repository under accession number GSE106095.

### Bioinformatic analysis of RNA-seq data

Sequencing files were demultiplexed using bcl2fastq (v2.19, Illumina), and quality determined using FastQC (https://www.bioinformatics.babraham.ac.uk/projects/fastqc/). A genomic reference index for *Drosophila melanogaster* was constructed with RSEM using the most recent genome build (BDGP release 6) and transcriptome annotation (Release 6.15) obtained from Flybase (www.flybase.org). Annotations used for lncRNAs were described in (Young et al. 2012). Paired-end reads were pseudo-aligned to the RSEM reference index using Salmon (Release 0.8.1) using the following parameters:

$ salmon quant --libType ISF –seqBias –gcBias –posBias –p 8 --numBootstraps 100. Gene-level counts were prepared for differential expression analysis with tximport as part of the Bioconductor package (Release 3.5). Feature length-scaled transcript counts per million reads (TPM) were calculated with tximport using the following command:

> tximport(files, type = “salmon”, countsFromAbundance = “lengthScaledTPM”, tx2gene = tx2gene)

Given the cell type heterogeneity between samples in this dataset, we used normalized counts instead of TPM or FPKM for more accurate inter-sample comparisons of gene abundance. We normalized gene-level counts via variance stabilizing transformation (File S1). Variance-stabilized transformed counts, principal component analysis (PCA), and differential expression was calculated using DESeq2 (Love et al. 2014) as part of the Bioconductor package (Release 3.5), using default parameters.

### PhyloCSF

*PhyloCSF* uses substitutions and codon frequencies in a genome alignment of 23 *Drosophilid* species to distinguish the evolutionary signature of selection for protein-coding function (Lin et al. 2011). For each transcript, PhyloCSF generates a score for the putative ORF with highest coding potential; transcripts with positive scores are more likely to be protein-coding. The candidate ORFs, their PhyloCSF scores, and other related information are included in File S5.

Briefly, local alignments used for PhyloCSF were extracted from the *23-Drosophilid* subset of the 27-way MULTIZ insect whole-genome alignments (Blanchette et al. 2004), downloaded from UCSC: http://hgdownload.soe.ucsc.edu/goldenPath/dm6/multiz27way/ (Tyner et al. 2017). PhyloCSF scores were computed using the 23flies parameters with the options “-f3 -- orf=ATGStop --allScores --bls”, which computes the score of every open reading frame (ORF) within the transcript that begins with ATG, is followed by a stop codon, and is at least the default length of 25 codons. Because CR44272 has no putative ORFs that long, we used “-- minCodons=19” to lower the threshold for that gene to the length of its longest putative ORF. We then selected the ORF in each transcript having the highest PhyloCSF score. The reported “ScorePerCodon” is the PhyloCSF score divided by the number of codons in the putative ORF. To identify potential cases in which one of the transcripts under consideration contains part of a coding ORF but the complete ORF is in an unidentified overlapping transcript, we also ran PhyloCSF using the --orf=StopStop3 option, with --minCodons=10, which looks for ORF fragments ending in a stop codon. However, that did not identify any plausible partial coding ORFs. The PhyloCSF track images in Figures 3 and S6 are overlays of the “Smoothed PhyloCSF” tracks in all three frames on the appropriate strand, from the PhyloCSF track hub in the UCSC genome browser, documented at: https://data.broadinstitute.org/compbio1/PhyloCSFtracks/trackHub/hub.DOC.html.

### Generation of coverage plots

The strand-specific and paired-end RNA-seq reads were mapped to the *Drosophila melanogaster* reference genome dm6 with the splicing-aware mapper STAR v2.5.3a (Dobin et al. 2013) using default parameters and a *Drosophila-specific* adjustment for maximum intron length and mate distance of 50kb. The resulting BAM files were filtered to include only uniquely mapping read pairs and then converted into strand-specific genome coverage tracks in BigWig format for visualization in the UCSC genome browser (Kent et al. 2010; Raney et al. 2014) using the program stranded-coverage (https://github.com/pmenzel/stranded-coverage) and wigToBigWig from the UCSC genome browser tools.

### Immunohistochemistry and FISH

Immunohistochemistry and RNA-FISH was performed as previously described (Kosman et al. 2004; Karaiskos et al. 2017). Primary and secondary antibodies used in this study are listed in Table S1. The procedure for probe synthesis is detailed in Supplementary Materials and Methods, and RNA probes are listed in Tables S3 and S4.

### Microscopy

Confocal stacks were imaged using the Leica SP8 equipped with 405 nm laser diode, white light laser (WLL), and hybrid detectors (HyD), with a 20x glycerol objective. For each field of view, 65-85 slices were acquired using ~AU=1 pinholes and taking care not to saturate signal. Appropriate slices were maximum intensity projected.

## Author contributions

A.L.M. designed and performed the majority of experiments and RNA-seq analysis, and adapted and optimized Fractionation-Seq for *Drosophila* embryos. S.W. performed several RNA-FISH experiments and provided critical revision of the manuscript. P.W. and A.L.M. developed and optimized the DIV-MARIS method for *Drosophila* embryos. P.W. generated the *IC::dsRed* and *VC::dsRed* fly strains and performed DIV-MARIS for 4-6h Ind samples. I.J. performed all PhyloCSF analyses. P.M. generated the coverage files. C.J.S. assisted with preliminary RNA-seq analyses. R.A. assisted with RNA-FISH experiments. I.M.M. acquired funding for Fractionation-Seq. R.P.Z. conceived the study, acquired funding for most experiments, and supervised the research. A.L.M. and R.P.Z. wrote the manuscript and all authors gave final approval for publication of the manuscript.

## Acknowledgements

We are grateful to John L. Rinn for insightful comments and extensive discussions, to Petar Glažar and Panagiotis Papavasileiou for considerable RNA-Seq troubleshooting and preliminary data analysis, Andrew Woehler and Joe Dragavon for assistance with confocal imaging, David Schechner for providing the cell fractionation protocol, Sara Ugowski and Claudia Kipar for maintaining the fly facility, and Sabrina Krüger and Agnieszka Klawiter for generating transgenic fly lines. Moreover, we are grateful to all members of the Zinzen Laboratory for in-depth discussions and technical assistance on experiments presented in this study.

A.L.M. and P.W. were supported by a generous grant from the Deutsche Forschungsgemeinschaft (DFG) Priority Programme SPP1738: Emerging roles of noncoding RNAs in nervous system development, plasticity, and disease. I.J. was supported by National Institutes of Health [HG004037] and GENCODE Wellcome Trust [U41 HG007234].

## Accession numbers

The NCBI Gene Expression Omnibus accession number for the RNA-seq data reported in this paper is GSE106095.

## References

Beckervordersandforth, R.M. et al., 2008. Subtypes of glial cells in the Drosophila embryonic ventral nerve cord as related to lineage and gene expression. Mechanisms of development, 125(5-6), pp.542–557.

Berger, C. et al., 2007. The commonly used marker ELAV is transiently expressed in neuroblasts and glial cells in the Drosophila embryonic CNS. Developmental dynamics : an official publication of the American Association of Anatomists, 236(12), pp.3562–3568.

Bier, E. & De Robertis, E.M., 2015. EMBRYO DEVELOPMENT. BMP gradients: A paradigm for morphogen-mediated developmental patterning. Science, 348(6242), pp.aaa5838–aaa5838.

Blanchette, M. et al., 2004. Aligning multiple genomic sequences with the threaded blockset aligner. Genome Research, 14(4), pp.708–715.

Bossing, T. et al., 1996. The embryonic central nervous system lineages of Drosophila melanogaster. I. Neuroblast lineages derived from the ventral half of the neuroectoderm. Developmental Biology, 179(1), pp.41–64.

Briggs, J.A. et al., 2015. Mechanisms of Long Non-coding RNAs in Mammalian Nervous System Development, Plasticity, Disease, and Evolution. Neuron, 88(5), pp.861–877.

Broadus, J. et al., 1995. New neuroblast markers and the origin of the aCC/pCC neurons in the Drosophila central nervous system. Mechanisms of development, 53(3), pp.393–402.

Brown, J.B. et al., 2014. Diversity and dynamics of the Drosophila transcriptome. Nature, 512(7515), pp.393–399.

Campos-Ortega, J.A., 1995. Genetic mechanisms of early neurogenesis in Drosophila melanogaster. Molecular neurobiology, 10(2-3), pp.75–89.

Chen, B. et al., 2016. Genome-wide identification and developmental expression profiling of long noncoding RNAs during Drosophila metamorphosis. Scientific reports, 6(1), p.23330.

Chen, L.-L., 2016. Linking Long Noncoding RNA Localization and Function. Trends in Biochemical Sciences, 41(9), pp.761–772.

Cowden, J. & Levine, M., 2003. Ventral dominance governs sequential patterns of gene expression across the dorsal-ventral axis of the neuroectoderm in the Drosophila embryo. Developmental Biology, 262(2), pp.335–349.

Crews, S.T., 2010. Axon-glial interactions at the Drosophila CNS midline. Cell Adhesion & Migration, 4(1), pp.1–5.

Dobin, A. et al., 2013. STAR: ultrafast universal RNA-seq aligner. Bioinformatics (Oxford, England), 29(1), pp.15–21.

Doe, C.Q., 1992. Molecular markers for identified neuroblasts and ganglion mother cells in the Drosophila central nervous system. Development, 116(4), pp.855–863.

Doe, C.Q., 2017. Temporal Patterning in the Drosophila CNS. Annual review of cell and developmental biology, 33(1), pp.219–240.

Goff, L.A. et al., 2015. Spatiotemporal expression and transcriptional perturbations by long noncoding RNAs in the mouse brain. Proceedings of the National Academy of Sciences, 112(22), pp.6855–6862.

Graveley, B.R. et al., 2011. The developmental transcriptome of Drosophila melanogaster. Nature, 471(7339), pp.473–479.

Hammonds, A.S. et al., 2013. Spatial expression of transcription factors in Drosophila embryonic organ development. Genome biology, 14(12), p.R140.

Heckscher, E.S. et al., 2014. Atlas-builder software and the eNeuro atlas: resources for developmental biology and neuroscience. Development, 141(12), pp.2524–2532.

Homem, C.C.F. & Knoblich, J.A., 2012. Drosophila neuroblasts: a model for stem cell biology. Development, 139(23), pp.4297–4310.

Hrvatin, S. et al., 2014. MARIS: Method for Analyzing RNA following Intracellular Sorting K. Aalto-Setala, ed. PLoS ONE, 9(3), pp.e89459–6.

Inagaki, S. et al., 2005. Identification and expression analysis of putative mRNA-like non-coding RNA in Drosophila. Genes to Cells, 10(12), pp.1163–1173.

Karaiskos, N. et al., 2017. The Drosophila embryo at single-cell transcriptome resolution. Science, 358(6360), pp.194–199.

Kent, W.J. et al., 2010. BigWig and BigBed: enabling browsing of large distributed datasets. Bioinformatics (Oxford, England), 26(17), pp.2204–2207.

Kosman, D. et al., 2004. Multiplex detection of RNA expression in Drosophila embryos. Science, 305(5685), pp.846–846.

Kunisch, M., Haenlin, M. & Campos-Ortega, J.A., 1994. Lateral inhibition mediated by the Drosophila neurogenic gene delta is enhanced by proneural proteins. Proceedings of the National Academy of Sciences, 91(21), pp.10139–10143.

Lai, E.C., 2004. Notch signaling: control of cell communication and cell fate. Development, 131(5), pp.965–973.

Landgraf, M. et al., 1997. The origin, location, and projections of the embryonic abdominal motorneurons of Drosophila. Journal of Neuroscience, 17(24), pp.9642–9655.

Landskron, L. et al., 2018. The asymmetrically segregating lncRNA cherub is required for transforming stem cells into malignant cells. eLife, 7, p.R106.

Lécuyer, E. et al., 2007. Global analysis of mRNA localization reveals a prominent role in organizing cellular architecture and function. Cell, 131(1), pp.174–187.

Li, M. & Liu, L., 2015. Neural functions of long noncoding RNAs in Drosophila. Journal of comparative physiology. A, Neuroethology, sensory, neural, and behavioral physiology, 201(9), pp.921–926.

Lin, M.F., Jungreis, I. & Kellis, M., 2011. PhyloCSF: a comparative genomics method to distinguish protein coding and non-coding regions. Bioinformatics (Oxford, England), 27(13), pp.i275–82.

Liu, S.J. et al., 2016. Single-cell analysis of long non-coding RNAs in the developing human neocortex. Genome biology, 17(1), p.67.

Love, M.I., Huber, W. & Anders, S., 2014. Moderated estimation of fold change and dispersion for RNA-seq data with DESeq2. Genome biology, 15(12), p.550.

McGovern, V.L. et al., 2003. A targeted gain of function screen in the embryonic CNS of Drosophila. Mechanisms of development, 120(10), pp.1193–1207.

Molyneaux, B.J. et al., 2015. DeCoN: Genome-wide Analysis of In Vivo Transcriptional Dynamics during Pyramidal Neuron Fate Selection in Neocortex. Neuron, 85(2), pp.275–288.

Ohlen, von T. & Doe, C.Q., 2000a. Convergence of dorsal, dpp, and egfr signaling pathways subdivides the drosophila neuroectoderm into three dorsal-ventral columns. Developmental Biology, 224(2), pp.362–372.

Ohlen, von T. & Doe, C.Q., 2000b. Convergence of dorsal, dpp, and egfr signaling pathways subdivides the drosophila neuroectoderm into three dorsal-ventral columns. Developmental Biology, 224(2), pp.362–372.

Picao-Osorio, J. et al., 2017. Pervasive Behavioral Effects of MicroRNA Regulation in Drosophila. Genetics, 206(3), pp.1535–1548.

Raney, B.J. et al., 2014. Track data hubs enable visualization of user-defined genome-wide annotations on the UCSC Genome Browser. Bioinformatics (Oxford, England), 30(7), pp.1003–1005.

Rickert, C. et al., 2011. Morphological characterization of the entire interneuron population reveals principles of neuromere organization in the ventral nerve cord of Drosophila. The Journal of neuroscience : the official journal of the Society for Neuroscience, 31(44), pp.15870–15883.

Rogers, W.A. et al., 2017. Uncoupling neurogenic gene networks in the Drosophila embryo. Genes & Development, 31(7), pp.634–638.

Sandler, J.E. & Stathopoulos, A., 2016. Stepwise Progression of Embryonic Patterning. Trends in Genetics, 32(7), pp.432–443.

Sauvageau, M. et al., 2013. Multiple knockout mouse models reveal lincRNAs are required for life and brain development. eLife, 2, p.e01749.

Schmidt, H. et al., 1997. The embryonic central nervous system lineages of Drosophila melanogaster. II. Neuroblast lineages derived from the dorsal part of the neuroectoderm. Developmental Biology, 189(2), pp.186–204.

Skeath, J.B. & Thor, S., 2003. Genetic control of Drosophila nerve cord development. Current Opinion in Neurobiology, 13(1), pp.8–15.

Skeath, J.B., Panganiban, G.F. & Carroll, S.B., 1994. The ventral nervous system defective gene controls proneural gene expression at two distinct steps during neuroblast formation in Drosophila. Development, 120(6), pp.1517–1524.

Sousa-Nunes, R., Cheng, L.Y. & Gould, A.P., 2010. Regulating neural proliferation in the Drosophila CNS. Current Opinion in Neurobiology, 20(1), pp.50–57.

Tomancak, P. et al., 2007. Global analysis of patterns of gene expression during Drosophila embryogenesis. Genome biology, 8(7), p.R145.

Tomancak, P. et al., 2002. Systematic determination of patterns of gene expression during Drosophila embryogenesis. Genome biology, 3(12), p.RESEARCH0088.

Tyner, C. et al., 2017. The UCSC Genome Browser database: 2017 update. Nucleic Acids Research, 45(D1), pp.D626–D634.

Weiss, J.B. et al., 1998. Dorsoventral patterning in the Drosophila central nervous system: the intermediate neuroblasts defective homeobox gene specifies intermediate column identity. Genes & Development, 12(22), pp.3591–3602.

Wen, K. et al., 2016. Critical roles of long noncoding RNAs in Drosophila spermatogenesis. Genome Research, 26(9), pp.1233–1244.

Wheeler, S.R. et al., 2006. Single-cell mapping of neural and glial gene expression in the developing Drosophila CNS midline cells. Developmental Biology, 294(2), pp.509–524.

Wheeler, S.R., Stagg, S.B. & Crews, S.T., 2009. MidExDB: a database of Drosophila CNS midline cell gene expression. BMC developmental biology, 9(1), p.56.

Wilk, R. et al., 2016. Diverse and pervasive subcellular distributions for both coding and long noncoding RNAs. Genes & Development, 30(5), pp.594–609.

Young, R.S. et al., 2012. Identification and Properties of 1,119 Candidate LincRNA Loci in the Drosophila melanogaster Genome. Genome Biology and Evolution, 4(4), pp.427–442.

Zhou, L. et al., 1995. Programmed cell death in the Drosophila central nervous system midline. Current Biology, 5(7), pp.784–790.

Santiago, C., Labrador, J.-P. & Bashaw, G.J., 2014. The homeodomain transcription factor Hb9 controls axon guidance in Drosophila through the regulation of Robo receptors. CellReports, 7(1), pp.153–165.

