## Supplemental Text and Figures for "A gene expression atlas of embryonic neurogenesis in *Drosophila* reveals complex spatiotemporal regulation of lncRNAs"

### 1. Supplementary Materials and Methods

#### *1.1 Fly strains and husbandry*

*Drosophila melanogaster* lines used in this study include

| | | |
|---|---|---|
| *yellow-white* | $y^1$ $w^{1118}$ | (source: BSC# 6598) |
| *IC*-GFP | $y^1$ $w^{1118}$ *; ; P{3xind_1.4-GFP}* | (this study) |
| *IC-dsRed* | $y^1$ $w^{1118}$ *; P{3xind_1.4-dsRed}* | (this study) |
| *VC-dsRed* | $y^1$ $w^{1118}$ *; ; P{2xvnd_743-dsRED}* | (Karaiskos et al. 2017) |
| *vnd-lexA/VC:FNLDD* | $y^1$ $w^{1118}$ *; M{3xvnd-lexA:Cit}*$^{attP\_ZH51C}$ *; P{vnd:FNLDD}* | (Krueger et al., unpublished) |
| *rho-lexA/VC:FNLDD* | $y^1$ $w^{1118}$ *; M{3xrho-lexA:Cit}*$^{attP\_ZH51C}$ *; P{vnd:FNLDD}* | (Krueger et al., unpublished) |

The *IC-GFP* and *IC-dsRed* lines were created by standard P-element transgenesis (Rubin & Spradling 1982) in a $y^1$ $w^{1118}$ background. The *ind* enhancer (*ind_1.4*) (Markstein et al. 2004) was cloned using primers (ngctagcgtcgacGCTTCAAAGCTCCGGGAAACG & nctcgagTCTGGGCCTTCGGTCCGAAAATG) flanked with NheI and SalI restriction sites (F primer) and with XhoI (R primer). PCR product was T/A cloned into pCRII-Duo and concatemerized using the compatibly cohesive sites XhoI and SalI. 3x *ind_1.4* constructs were directionally subcloned into the P-element vectors pH-Stinger or pRed-HStinger (Barolo et al. 2004) using NheI and XhoI.

Fly stocks were maintained at 25°C, ~60% relative humidity on standard fly food with 12 hr light/dark cycles according to standard procedures. 2-hr embryo collections were done on apple juice agar plates with yeast paste after 3 x 1hr pre-lays in the morning, embryos were then aged for an appropriate amount of time at 25°C and ~60% relative humidity before dechorionation.

#### *1.2 DIV-SortSeq*

This protocol incorporates portions from MARIS (Hrvatin et al. 2014). All steps after dechorionation were performed on ice, using ice-cold DEPC-treated solutions. Primary and secondary antibodies used in this study are listed in Table S1. RNase-free BSA was obtained from Gemini Bioproducts and is critical for isolation of high-quality RNA.

Embryos were collected, dechorionated, and dissociated into single-cell suspension in 15ml PBS, pH 7.5 using 8-12 strokes with the loose pestle of a glass dounce homogenizer. Large debris was removed via filtration with 2 x 90°-rotated sheets of Miracloth into a 15ml conical vial, followed by centrifugation at 40x*g*, 4°C, 3 min. The supernatant was transferred to a 15ml conical vial, cells were pelleted at 1000x*g*, 4°C, 3 min, washed with 1ml PBS, and fixed with 4% formaldehyde at 4°C, 15min. Cross-linking was stopped with Quench Buffer (2.5M Glycine in PBS). Fixed cells were washed twice with 1ml ice-cold PBS, and stored at 4°C overnight in 1ml RNA*later*™ (Thermo, AM7020). Cells were rehydrated via dilution in 9ml ice-cold PBS, centrifugation at 1000 x*g*, 10min, and washed twice in 1ml ice-cold PBS. Fixed cell suspensions were immunostained under RNase-free conditions with primary antibodies with agitation at

4°C, 1.5-2h in 250μl Stain Buffer (1% BSA (w/v), 0.1% saponin, 1:200 RNase inhibitor in RNase-free PBS). After washing 3x 5 min in 1ml Wash Buffer (0.2% BSA, 0.1% saponin in PBS), cells were incubated with conjugated secondary antibodies with agitation at 4°C, 45 min-1h in 250μl Stain Buffer. After washing 3x 5 min in 1ml Wash Buffer, cells were resuspended in 1ml Sort Buffer (0.5% BSA, 2mM EDTA, 1:500 RNase inhibitor in PBS) and filtered with a 70μm cell strainer. Filtered cells were subjected to FACS purification on the FACSAria II (BD Biosciences) and sorted cells collected in Collection Buffer (2% BSA, 1:100 RNase inhibitor in PBS).

FACS-purified cells were pelleted and resuspended in 200μl Digestion Buffer (5M NaCl, 1M Tris-HCl pH 8.0, 200mM EDTA, 10% SDS, 3.2U Proteinase K, 1:100 RNase inhibitor) and incubated at 50°C, 15min (Proteinase K digestion) followed by 80°C, 15min (reversal of formaldehyde cross-links). Samples were transferred to ice and resuspended in 600μl TRIzol™ LS Reagent (Thermo, 10296028). RNA was isolated using the DirectZol™ RNA MicroPrep Kit (Zymo Research, R2060) according to the manufacturer's instructions. RNA concentration was measured using the Qubit™ RNA HS Assay kit (Thermo, Q32852) and RNA quality determined using the Agilent RNA 6000 Pico Kit (Agilent, 5067-1513). All RNA-seq libraries were constructed using the NuGEN Ovation *Drosophila* RNA-Seq System with 10 ng – 100 ng total RNA input. Library concentration was quantified using the Qubit™ dsDNA HS Assay (Thermo, Q32854) and quality was determined on a BioAnalyzer™ using Agilent High Sensitivity DNA Kits (Agilent, 5067-4626). All libraries were sequenced on the Illumina HiSeq4000 at a mean depth of 62.5 million 75bp paired-end reads per sample. RNA-seq datasets generated for this study are detailed in Tables S5 and S6. A detailed, step-wise protocol is available upon request.

##### *1.3 Nuclear-cytoplasmic fractionation*

The cell fractionation procedure incorporates portions from MARIS (Hrvatin et al. 2014). All steps were performed on ice, using ice-cold DEPC-treated solutions, and all centrifugation steps were performed at 4˚C. Embryos were processed to single-cell suspension as described above for *DIV*-SortSeq, then pelleted and resuspended in Cyto Extract Buffer (20mM Tris pH 7.6, 0.1mM EDTA, 2mM $MgCl_2$). After hypotonic swelling, cells were gently lysed by addition of 0.6% CHAPS for isolation of the cytoplasmic fraction. Nuclei were pelleted at 500x*g* for 5min, and the supernatant retained, and an appropriate volume of TRIzol™ LS Reagent was added (cytoplasmic fraction).

Nuclei were washed with Nuclei Wash Buffer (20mM Tris pH 7.6, 0.1mM EDTA, 2mM $MgCl_2$, 0.6% CHAPS), resuspended in Nuclei Resuspension Buffer (10mM Tris, pH 7.6, 150mM NaCl, 0.15% NP-40) and pelleted at 12,000x*g*, 10min in Sucrose Buffer (10mM Tris, pH 7.5, 150mM NaCl, 24% sucrose). After washing (1mM EDTA in PBS), nuclei pellet was resuspended in TRIzol™ Reagent. RNA was isolated and concentration and quality determined as described for above for *DIV*-SortSeq.

*1.4 Quantitative RT-PCR (qPCR)*

50ng of total RNA was reverse-transcribed with the QuantiTect Reverse Transcription kit (Qiagen, 205310) in a total volume of 20*μ*l, according to the manufacturer’s instructions. For each gene, 0.2*μ*l cDNA was used for input into qPCR using SensiFAST™ SYBR® No-ROX Kit (Bioline, 98020) and 5*μ*M forward and reverse primers in a total volume of 20*μ*l. qPCR primer sequences are listed in Table S2. qPCR thermal cycling and fluorescent data acquisition was performed using the BioRad CFX96 Touch™ Real-Time PCR Detection System. Expression fold changes were calculated via the $\Delta\Delta C_T$ method (Vandesompele et al. 2002; Schmittgen & Livak 2008), normalized to the mean $C_T$ of two reference genes: *α-tubulin* and *actin 42A*.

*1.5 RNA probe design and synthesis for RNA in situ hybridization.*

Where available, cDNA clones were obtained from the *Drosophila* Gene Collection or *Drosophila* Genomics Resource Center (Stapleton et al. 2002), detailed in Table S3. Constructs were linearized via restriction digestion, and subjected to *in vitro* transcription using appropriate RNA Polymerases (Roche), using ribonucleotide mixtures containing dUTP-DIG, -FITC, or -Biotin (Roche).

Where cDNA clones were not available, PCR primers were designed to amplify a region within the transcribed locus from genomic DNA. Primer sequences are detailed in Table S4. A T7 promoter sequence appended to the reverse primer allowed *in vitro* transcription directly from the PCR product. After template digest by DNaseI, RNA probes were sheared at 65°C for 3-20 min in carbonation buffer (120mM $Na_2CO_3$, 80mM $NaHCO_3$, pH 10.2), length of carbonation depended on probe length. RNA probes were precipitated at -20°C overnight and resuspended in Hyb-A Buffer (50% formamide, 5X SSC, 0.1% Tween-20).

#### 2. Supplementary Tables

*Table S1: Primary and secondary antibodies used in this study*

| *Target* | *Supplier* | *Catalog #* | *Application* | *Antibody Type* | *Conjugate* | *Dilution* |
|---|---|---|---|---|---|---|
| *RFP* | Thermo | 710530 | *Primary* | Rabbit polyclonal | N/A | 1:500 |
| *GFP* | Thermo | G10362 | *Primary* | Rabbit polyclonal | N/A | 1:500 |
| *Pros* | DSHB | MR1A | *Primary* | Mouse monoclonal | N/A | 1:20 |
| *Elav* | DSHB | 9F8A9 | *Primary* | Mouse monoclonal | N/A | 1:500 |
| *Repo* | DSHB | 8D12 | *Primary* | Mouse monoclonal | N/A | 1:20 |
| *Rabbit IgG* | Thermo | A21428 | *Secondary* | Goat polyclonal | *Alexa Fluor 555* | 1:500 |
| *Mouse IgG* | Thermo | A32727 | *Secondary* | Goat polyclonal | *Alexa Fluor 555* | 1:500 |

*Table S2: qPCR primer sequences*

| *Target* | *Forward primer* | *Reverse primer* | *Size (bp)* |
|---|---|---|---|
| *α-Tubulin* | TGTCGCGTGTGAAACACTTC | AGCAGGCGTTTCCAATCTG | 585 |
| *Actin 42A* | GCGTCGGTCAATTCAATCTT | AAGCTGCAACCTCTTCGTCA | 292 |
| *Prospero* | CGGCATGGCTCCTACTTCTT | TAGCGCACCCAGAAGAACAT | 78 |
| *Worniu* | ATGGATAAACTCAAGTACAGCCG | AAGTCCACTGGTCCTTCATCA | 107 |
| *Elav* | ACGCTCCTGCCACAGAAAAA | CGTCGCCGTATTTCGCTC | 211 |
| *Lim3* | GATGGAGGATCGTAAGCTGATCT | GTAGGCCGTTTTCAGGGTCTC | 154 |
| *Repo* | CTCCGCCAAGTAGTTCCTCC | AGGCAGTAAAGGTGGTTCTCG | 216 |
| *Gcm* | ACAAGGCCAGAAGGAAGCAG | CAAGCCTGGATTTCCAAGCGA | 76 |

*Table S3: List of commercially-available cDNA clones for RNA probe synthesis*

| *RNA target* | *DGRC Reference #* |
|---|---|
| *ind* | RT01026 |
| *vnd* | PCSP6029 |
| *pros* | LD37627 |
| *elav* | LD33076-IR |
| *repo* | GH05443-dg |
| *CR30009* | RE30084 |
| *CR32730* | RE54940 |
| *CR32111* | RE52337 |

*Table S4: PCR primer sequences for RNA probe synthesis*

| *Target* | *Forward primer* | *T7 promoter + Reverse primer* | *Size (bp)* |
|---|---|---|---|
| *CR46003* | TGTGTCGCACAGGATGTGT | TAATACGACTCACTATAGGTGCTGGCGGGGAAATTATGT | 908 |
| *cherub* | CGAGGAACCTTCGGTGCATA | TAATACGACTCACTATAGGGCTTGGGTGATTTCGAGGGA | 1511 |
| *CR44024* | GTGTCGTGTCGGGTAAGTGT | TAATACGACTCACTATAGGAAGTGGCCTGTCTCAGAACG | 1268 |
| *CR32111* | GTATGCGCTCGAACTCGGTAA | TAATACGACTCACTATAGGGCCGGCATGAGCAAACACAAA | 1232 |

*Table S5: Summary of RNA-seq datasets – DIV-SortSeq*

| ***Sample name*** | ***Fly line*** | ***Target protein*** | ***Cell Type*** | ***Enriched/depleted*** | ***Time point*** | ***Replicate*** | ***# reads (M)*** |
|---|---|---|---|---|---|---|---|
| *4-6h_Ind-neg_1* | *IC*-dsRed | dsRed | Intermediate column | Depleted | 4-6h | 1 | 89.9 |
| *4-6h_Ind-neg_2* | *IC*-dsRed | dsRed | Intermediate column | Depleted | 4-6h | 2 | 140.1 |
| *4-6h_Ind-pos_1* | *IC*-dsRed | dsRed | Intermediate column | Enriched | 4-6h | 1 | 76.2 |
| *4-6h_Ind-pos_2* | *IC*-dsRed | dsRed | Intermediate column | Enriched | 4-6h | 2 | 81.6 |
| *4-6h_Vnd-neg_1* | *VC*-dsRed | dsRed | Ventral column | Depleted | 4-6h | 1 | 62.6 |
| *4-6h_Vnd-neg_2* | *VC*-dsRed | dsRed | Ventral column | Depleted | 4-6h | 2 | 64.3 |
| *4-6h_Vnd-pos_1* | *VC*-dsRed | dsRed | Ventral column | Enriched | 4-6h | 1 | 61.4 |
| *4-6h_Vnd-pos_2* | *VC*-dsRed | dsRed | Ventral column | Enriched | 4-6h | 2 | 65.6 |
| *4-6h_Pros-neg_1* | *vnd*-lexA/VC:FNLDD | Prospero | Neuroblasts | Depleted | 4-6h | 1 | 59.8 |
| *4-6h_Pros-neg_2* | *vnd*-lexA/VC:FNLDD | Prospero | Neuroblasts | Depleted | 4-6h | 2 | 79.1 |
| *4-6h_Pros-pos_1* | *vnd*-lexA/VC:FNLDD | Prospero | Neuroblasts | Enriched | 4-6h | 1 | 61.9 |
| *4-6h_Pros-pos_2* | *vnd*-lexA/VC:FNLDD | Prospero | Neuroblasts | Enriched | 4-6h | 2 | 59.1 |
| *6-8h_Ind-neg_1* | *IC*-GFP | GFP | Intermediate column | Depleted | 6-8h | 1 | 61 |
| *6-8h_Ind-neg_2* | *IC*-GFP | GFP | Intermediate column | Depleted | 6-8h | 2 | 59.9 |
| *6-8h_Ind-pos_1* | *IC*-GFP | GFP | Intermediate column | Enriched | 6-8h | 1 | 57 |
| *6-8h_Ind-pos_2* | *IC*-GFP | GFP | Intermediate column | Enriched | 6-8h | 2 | 70.9 |
| *6-8h_Vnd-neg_1* | *VC*-dsRed | dsRed | Ventral column | Depleted | 6-8h | 1 | 73.2 |
| *6-8h_Vnd-neg_2* | *VC*-dsRed | dsRed | Ventral column | Depleted | 6-8h | 2 | 61 |
| *6-8h_Vnd-pos_1* | *VC*-dsRed | dsRed | Ventral column | Enriched | 6-8h | 1 | 61.6 |
| *6-8h_Vnd-pos_2* | *VC*-dsRed | dsRed | Ventral column | Enriched | 6-8h | 2 | 73.9 |
| *6-8h_Pros-neg_1* | *vnd*-lexA/VC:FNLDD | Prospero | Neuroblasts | Depleted | 6-8h | 1 | 64.5 |
| *6-8h_Pros-neg_2* | *vnd*-lexA/VC:FNLDD | Prospero | Neuroblasts | Depleted | 6-8h | 2 | 76.6 |
| *6-8h_Pros-pos_1* | *vnd*-lexA/VC:FNLDD | Prospero | Neuroblasts | Enriched | 6-8h | 1 | 54.3 |
| *6-8h_Pros-pos_2* | *vnd*-lexA/VC:FNLDD | Prospero | Neuroblasts | Enriched | 6-8h | 2 | 58.8 |
| *6-8h_Elav-neg_1* | *IC*-GFP | Elav | Neurons | Depleted | 6-8h | 1 | 62.2 |
| *6-8h_Elav-neg_2* | *yw* | Elav | Neurons | Depleted | 6-8h | 2 | 57.2 |
| *6-8h_Elav-pos_1* | *IC*-GFP | Elav | Neurons | Enriched | 6-8h | 1 | 53.8 |
| *6-8h_Elav-pos_2* | *yw* | Elav | Neurons | Enriched | 6-8h | 2 | 70.4 |
| *6-8h_Repo-neg_1* | *IC*-GFP | Repo | Glia | Depleted | 6-8h | 1 | 59.6 |
| *6-8h_Repo-neg_2* | *IC*-GFP | Repo | Glia | Depleted | 6-8h | 2 | 71.4 |
| *6-8h_Repo_pos_1* | *IC*-GFP | Repo | Glia | Enriched | 6-8h | 1 | 66.6 |
| *6-8h_Repo_pos_2* | *IC*-GFP | Repo | Glia | Enriched | 6-8h | 2 | 56.8 |
| *8-10h_Elav-neg_1* | *y w* | Elav | Neurons | Depleted | 8-10h | 1 | 74.5 |
| *8-10h_Elav-neg_2* | *y w* | Elav | Neurons | Depleted | 8-10h | 2 | 44.1 |
| *8-10h_Elav-pos_1* | *y w* | Elav | Neurons | Enriched | 8-10h | 1 | 69.9 |
| *8-10h_Elav-pos_2* | *y w* | Elav | Neurons | Enriched | 8-10h | 2 | 62.9 |
| *8-10h_Repo-neg_1* | *vnd*-lexA/VC:FNLDD | Repo | Glia | Depleted | 8-10h | 1 | 61.5 |
| *8-10h_Repo-neg_2* | *vnd*-lexA/VC:FNLDD | Repo | Glia | Depleted | 8-10h | 2 | 60.9 |
| *8-10h_Repo-pos_1* | *vnd*-lexA/VC:FNLDD | Repo | Glia | Enriched | 8-10h | 1 | 61.4 |

| | | | | | | | |
|---|---|---|---|---|---|---|---|
| *8-10h_Repo-pos_2* | *vnd-*lexA/VC:FNLDD | Repo | Glia | Enriched | 8-10h | 2 | 65.6 |
| *18-22h_Elav-neg_1* | *vnd-*lexA/VC:FNLDD | Elav | Neurons | Depleted | 18-22h | 1 | 68.6 |
| *18-22h_Elav-neg_2* | *vnd-*lexA/VC:FNLDD | Elav | Neurons | Depleted | 18-22h | 2 | 61.9 |
| *18-22h_Elav-pos_1* | *vnd-*lexA/VC:FNLDD | Elav | Neurons | Enriched | 18-22h | 1 | 66 |
| *18-22h_Elav-pos_2* | *vnd-*lexA/VC:FNLDD | Elav | Neurons | Enriched | 18-22h | 2 | 63.9 |
| *18-22h_Repo-neg_1* | *vnd-*lexA/VC:FNLDD | Repo | Glia | Depleted | 18-22h | 1 | 58.3 |
| *18-22h_Repo-neg_2* | *vnd-*lexA/VC:FNLDD | Repo | Glia | Depleted | 18-22h | 2 | 63.4 |
| *18-22h_Repo-pos_1* | *vnd-*lexA/VC:FNLDD | Repo | Glia | Enriched | 18-22h | 1 | 70.5 |
| *18-22h_Repo-pos_2* | *vnd-*lexA/VC:FNLDD | Repo | Glia | Enriched | 18-22h | 2 | 75.9 |

*Table S6: Summary of RNA-seq datasets – Fractionation-Seq*

| ***Sample name*** | ***Fly line*** | ***Fraction*** | ***Time point*** | ***Replicate*** | ***Number of reads (M)*** |
|---|---|---|---|---|---|
| *Cyto_18-22h_1* | *rho*-lexA/VC:FNLDD | Cytoplasmic | 18-22h | 1 | 163.2 |
| *Cyto_18-22h_2* | *rho*-lexA/VC:FNLDD | Cytoplasmic | 18-22h | 2 | 179.1 |
| *Cyto_6-8h_1* | *rho*-lexA/VC:FNLDD | Cytoplasmic | 6-8h | 1 | 189.3 |
| *Cyto_6-8h_2* | *rho*-lexA/VC:FNLDD | Cytoplasmic | 6-8h | 2 | 179.3 |
| *Nuc_18-22h_1* | *rho*-lexA/VC:FNLDD | Nuclear | 18-22h | 1 | 203 |
| *Nuc_18-22h_2* | *rho*-lexA/VC:FNLDD | Nuclear | 18-22h | 2 | 176.1 |
| *Nuc_6-8h_1* | *rho*-lexA/VC:FNLDD | Nuclear | 6-8h | 1 | 193.9 |
| *Nuc_6-8h_2* | *rho*-lexA/VC:FNLDD | Nuclear | 6-8h | 2 | 193.7 |
| *Whole_18-22h_1* | *rho*-lexA/VC:FNLDD | Whole embryo | 18-22h | 1 | 183.2 |
| *Whole_18-22h_2* | *rho*-lexA/VC:FNLDD | Whole embryo | 18-22h | 2 | 180.7 |
| *Whole_6-8h_1* | *rho*-lexA/VC:FNLDD | Whole embryo | 6-8h | 1 | 197.4 |
| *Whole_6-8h_2* | *rho*-lexA/VC:FNLDD | Whole embryo | 6-8h | 2 | 196.1 |

#### *2.1 Table S7: All protein-coding ‘computed genes’ correlated (r > 0.9) with neurogenic marker genes in DIV-SortSeq expression data*

| *Flybase ID* | *Annotation ID* |
|---|---|
| **FBgn0036725** | CG18265 |
| **FBgn0034009** | CG8155 |
| **FBgn0036008** | CG3408 |
| **FBgn0030017** | CG2278 |
| **FBgn0264449** | CG43867 |
| **FBgn0051235** | CG31235 |
| **FBgn0037166** | CG11426 |
| **FBgn0032752** | CG10702 |
| **FBgn0038114** | CG11670 |
| **FBgn0037206** | CG12768 |
| **FBgn0032485** | CG9426 |
| **FBgn0038321** | CG6218 |
| **FBgn0038720** | CG6231 |
| **FBgn0033283** | CG11635 |
| **FBgn0043806** | CG32032 |
| **FBgn0033287** | CG8701 |
| **FBgn0031540** | CG3238 |
| **FBgn0031961** | CG7102 |
| **FBgn0040984** | CG4440 |
| **FBgn0026876** | CG11403 |
| **FBgn0035689** | CG7376 |
| **FBgn0025388** | CG12179 |
| **FBgn0036503** | CG13454 |
| **FBgn0040809** | CG13465 |
| **FBgn0032489** | CG15480 |
| **FBgn0041702** | CG15107 |
| **FBgn0034403** | CG18190 |
| **FBgn0031070** | CG12702 |
| **FBgn0250754** | CG42232 |
| **FBgn0261538** | CG42662 |
| **FBgn0034514** | CG13427 |
| **FBgn0036670** | CG13029 |
| **FBgn0031252** | CG13690 |
| **FBgn0040346** | CG3704 |
| **FBgn0039013** | CG4813 |
| **FBgn0037633** | CG9839 |
| **FBgn0028506** | CG4455 |
| **FBgn0037844** | CG4570 |
| **FBgn0037924** | CG14712 |
| **FBgn0035464** | CG12006 |
| **FBgn0027602** | CG8611 |
| **FBgn0037051** | CG10565 |
| **FBgn0052756** | CG32756 |
| **FBgn0036710** | CG6479 |
| **FBgn0039735** | CG7911 |
| **FBgn0023515** | CG14814 |
| **FBgn0033160** | CG11107 |
| **FBgn0033169** | CG11123 |
| **FBgn0029672** | CG2875 |
| **FBgn0025633** | CG13366 |
| **FBgn0050183** | CG30183 |
| **FBgn0030768** | CG9723 |
| **FBgn0025626** | CG4281 |
| **FBgn0035213** | CG2199 |
| **FBgn0031403** | CG15387 |
| **FBgn0043456** | CG4747 |
| **FBgn0052428** | CG32428 |
| **FBgn0031062** | CG14230 |
| **FBgn0035315** | CG8960 |
| **FBgn0032050** | CG13096 |
| **FBgn0038552** | CG18012 |
| **FBgn0031764** | CG9107 |
| **FBgn0035878** | CG7182 |
| **FBgn0034073** | CG8414 |
| **FBgn0033766** | CG8771 |
| **FBgn0037958** | CG6962 |
| **FBgn0034447** | CG7744 |
| **FBgn0035842** | CG7504 |
| **FBgn0038272** | CG7265 |
| **FBgn0039544** | CG12877 |
| **FBgn0033615** | CG7741 |
| **FBgn0030122** | CG16892 |
| **FBgn0051365** | CG31365 |
| **FBgn0052318** | CG32318 |
| **FBgn0039566** | CG4849 |
| **FBgn0037746** | CG8478 |
| **FBgn0030813** | CG4949 |
| **FBgn0029825** | CG12728 |
| **FBgn0038768** | CG4936 |
| **FBgn0037149** | CG14561 |
| **FBgn0037372** | CG2091 |
| **FBgn0030915** | CG6179 |
| **FBgn0027514** | CG1024 |
| **FBgn0031947** | CG7154 |
| **FBgn0030660** | CG8097 |
| **FBgn0036565** | CG5235 |
| **FBgn0024364** | CG11417 |
| **FBgn0036994** | CG5199 |
| **FBgn0033527** | CG11777 |
| **FBgn0039743** | CG7946 |
| **FBgn0262719** | CG43163 |
| **FBgn0036886** | CG9300 |
| **FBgn0034264** | CG10933 |
| **FBgn0025627** | CG4194 |
| **FBgn0032682** | CG10176 |
| **FBgn0031001** | CG7884 |
| **FBgn0037213** | CG12581 |
| **FBgn0038660** | CG14291 |
| **FBgn0034933** | CG3735 |
| **FBgn0030855** | CG5800 |
| **FBgn0035414** | CG14965 |
| **FBgn0036214** | CG7264 |
| **FBgn0030317** | CG1561 |
| **FBgn0036660** | CG13025 |
| **FBgn0031597** | CG17612 |
| **FBgn0036483** | CG12316 |
| **FBgn0032512** | CG9305 |
| **FBgn0037644** | CG11964 |
| **FBgn0040385** | CG12496 |
| **FBgn0033802** | CG17724 |
| **FBgn0039733** | CG11504 |
| **FBgn0037504** | CG1142 |
| **FBgn0036522** | CG7372 |
| **FBgn0035402** | CG12082 |
| **FBgn0031769** | CG9135 |
| **FBgn0038551** | CG7357 |
| **FBgn0032348** | CG4751 |
| **FBgn0035987** | CG3689 |
| **FBgn0037918** | CG6791 |
| **FBgn0035872** | CG7185 |
| **FBgn0031529** | CG9662 |
| **FBgn0031492** | CG3542 |
| **FBgn0086855** | CG17078 |
| **FBgn0033990** | CG10265 |
| **FBgn0032454** | CG5787 |
| **FBgn0037622** | CG8202 |
| **FBgn0032751** | CG17343 |
| **FBgn0266917** | CG16941 |
| **FBgn0034114** | CG4282 |
| **FBgn0034750** | CG3732 |
| **FBgn0030738** | CG9915 |
| **FBgn0035235** | CG7879 |
| **FBgn0028474** | CG4119 |
| **FBgn0085451** | CG34422 |
| **FBgn0030293** | CG1737 |
| **FBgn0050020** | CG30020 |
| **FBgn0029941** | CG1677 |
| **FBgn0033021** | CG10417 |
| **FBgn0027503** | CG11970 |
| **FBgn0032388** | CG6686 |
| **FBgn0038546** | CG7379 |
| **FBgn0035481** | CG12605 |
| **FBgn0260451** | CG14042 |
| **FBgn0035677** | CG13293 |
| **FBgn0036202** | CG6024 |
| **FBgn0035643** | CG13287 |
| **FBgn0034184** | CG9646 |
| **FBgn0052105** | CG32105 |
| **FBgn0031257** | CG4133 |
| **FBgn0037050** | CG10566 |
| **FBgn0025712** | CG13920 |
| **FBgn0027550** | CG6495 |
| **FBgn0031762** | CG9098 |
| **FBgn0035246** | CG13928 |
| **FBgn0039808** | CG12071 |
| **FBgn0030508** | CG15760 |
| **FBgn0033960** | CG10151 |
| **FBgn0030596** | CG12398 |
| **FBgn0052085** | CG32085 |
| **FBgn0030012** | CG18262 |
| **FBgn0030223** | CG2111 |
| **FBgn0051030** | CG31030 |
| **FBgn0035903** | CG6765 |
| **FBgn0030595** | CG14406 |
| **FBgn0033983** | CG10253 |
| **FBgn0031955** | CG14535 |
| **FBgn0033497** | CG12912 |
| **FBgn0034154** | CG5267 |
| **FBgn0033872** | CG6329 |
| **FBgn0030588** | CG9521 |
| **FBgn0085382** | CG34353 |
| **FBgn0085218** | CG34189 |
| **FBgn0030594** | CG9509 |
| **FBgn0034459** | CG16716 |
| **FBgn0259823** | CG42404 |
| **FBgn0032800** | CG10137 |
| **FBgn0038926** | CG13409 |
| **FBgn0039064** | CG4467 |
| **FBgn0259163** | CG42268 |
| **FBgn0259994** | CG42492 |
| **FBgn0039030** | CG6660 |
| **FBgn0036579** | CG5027 |
| **FBgn0029708** | CG3556 |
| **FBgn0030586** | CG12539 |
| **FBgn0038596** | CG14312 |
| **FBgn0263072** | CG43347 |
| **FBgn0037525** | CG17816 |
| **FBgn0030742** | CG9919 |
| **FBgn0265084** | CG44195 |
| **FBgn0030592** | CG9514 |
| **FBgn0262476** | CG43066 |
| **FBgn0035033** | CG3548 |
| **FBgn0040351** | CG11638 |
| **FBgn0028647** | CG11902 |
| **FBgn0031313** | CG5080 |
| **FBgn0039915** | CG1732 |
| **FBgn0029896** | CG3168 |
| **FBgn0039024** | CG4721 |
| **FBgn0032897** | CG9336 |
| **FBgn0034128** | CG4409 |
| **FBgn0034417** | CG15117 |
| **FBgn0031816** | CG16947 |
| **FBgn0030261** | CG15203 |
| **FBgn0033446** | CG1648 |
| **FBgn0052354** | CG32354 |
| **FBgn0031627** | CG15630 |
| **FBgn0036927** | CG7433 |
| **FBgn0034618** | CG9485 |
| **FBgn0083972** | CG34136 |
| **FBgn0031589** | CG3714 |
| **FBgn0036760** | CG5567 |
| **FBgn0032899** | CG9338 |

#### 3. Supplementary Notes

##### *3.1 Column interpretation for File S4: PhyloCSF*

| | |
|---|---|
| Intervals | Intervals of the input transcript |
| Strand | Strand |
| TranscriptName | Name from the input bed file |
| TrUCSCview | Link to show the entire transcript in UCSC browser |
| ORFintervals | Intervals of the ORF (not including the stop codon) |
| ORFstart | 0-based transcript coordinate of first base of ORF |
| ORFend | 0-based transcript coordinate of last base of ORF |
| NumCodons | Number of codons in the ORF |
| PhyloCSF | Raw PhyloCSF score of the ORF |
| RelBL | Fraction of branch length of the phylogenetic tree spanned by species present in the alignment of this ORF |
| ScorePerCodon | PhyloCSF divided by NumCodons |
| PhyloCSFPsi | Length adjusted score, a log likelihood, in decibans |
| Pval | Probability a region of this length, none-of-which has ever been coding, has this score or higher |
| CorrectedPvalTr | p-val with Holm-Bonferroni correction for number of ORFs in this transcript |
| CorrectedPvalAll | p-val with Holm-Bonferroni correction for total number of ORFs |
| FDR | Benjamini & Hochberg false discovery rate. |
| LocalFDR | Local FDR (Efron et al. 2001). |
| AntiScorePerCodon | Score on opposite strand in frame that shares 3rd codon position |
| ScoreDiff | ScorePerCodon - AntiScorePerCodon |
| GC | GC content of ORF |
| CpGratio | Number of CpGs in ORF divided by the expected number based on C and G content |
| CodAlignView | Link to view ORF alignment in CodAlignView, the Codon Alignment Viewer, with 10-codon context on each side |
| OrfUCSCview | Link to show the ORF in UCSC browser |

#### 4. Supplementary Figures

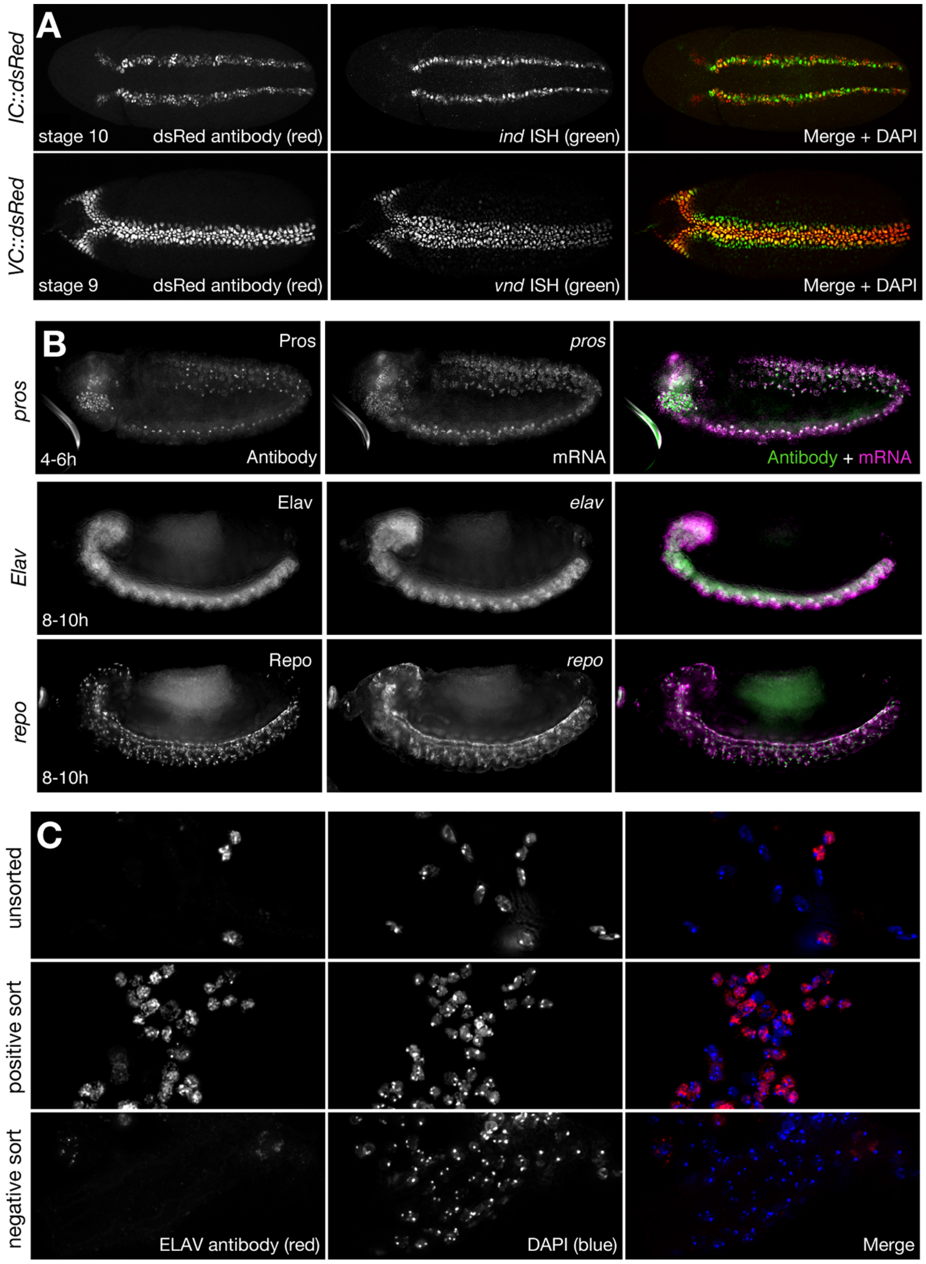

**Fig. S1: Neurogenic cell populations can be faithfully marked and purified**
(A) Visualization of fluorescent reporter expression in *VC-dsRed* (top panel) and *IC-dsRed* (bottom panel) transgenic embryos. Tissue reporter shown on left and in red by antibody stain, expression of the tissue marker is shown in the middle and in green by *in situ* hybridization. Shown are whole mount embryos, ventral views anterior left. (B) Multiplex whole mount immunohistochemistry (green) and RNA-FISH (magenta) show faithfulness of the antibody sorting markers for neuroblasts (Pros; top panel) at 4-6h, and neurons (Elav; middle panel) and glia (Repo; bottom panel) at 8-10h. (C) Immunohistochemical staining of endogenous Elav protein in cells of dissociated embryos pre- and post-FACS.

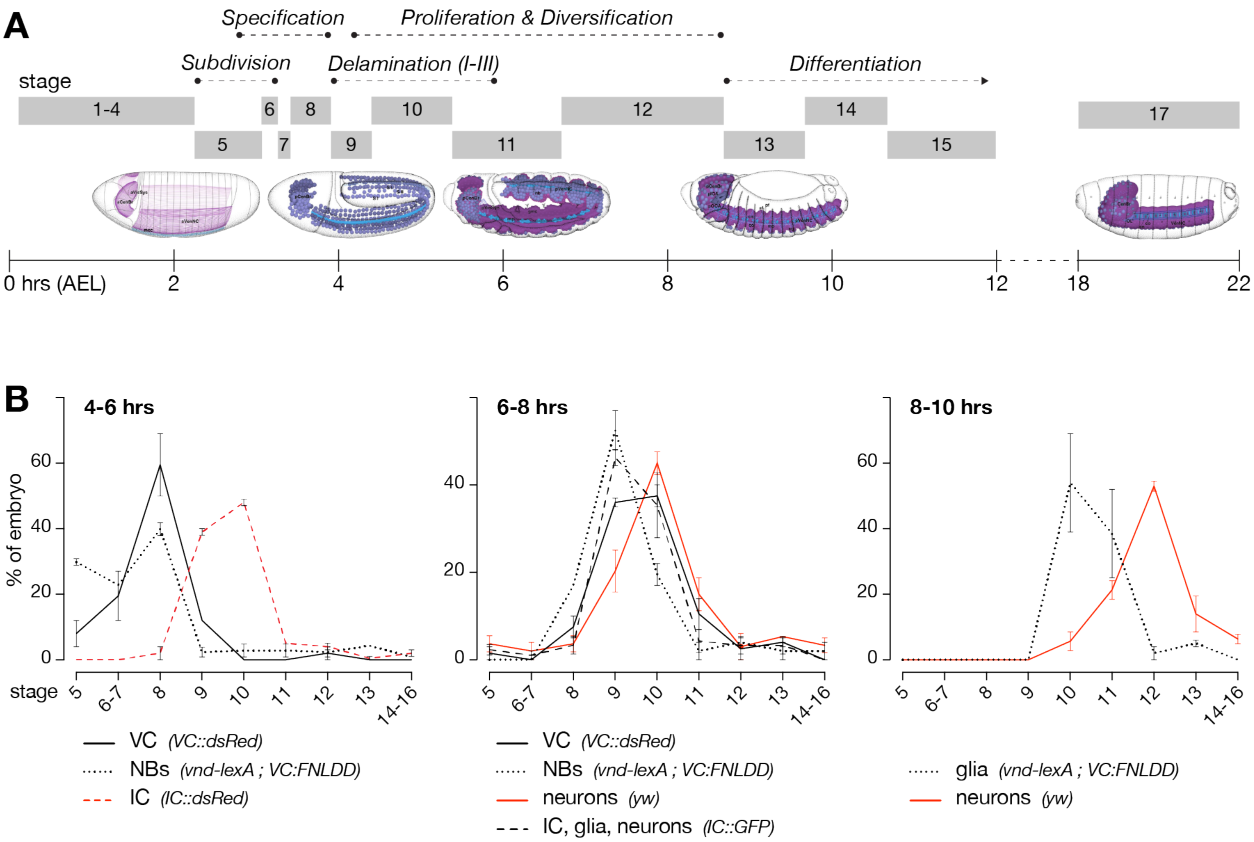

**Fig. S2: Timed embryo collections encompass major neurogenic events**
(A) Overview of *Drosophila* embryonic neurogenesis, timing and stages. Embryo images show stages of neurogenesis schematically and were obtained from the *Atlas of Drosophila Development* by Volker Hartstein (CSHL Press, 1992, used with permission). (B) Staging of representative samples corresponding to timed embryo collections (n = 2-3 collections per line). Staging according to (Campos-Ortega & Hartenstein 1985). Cell type markers targeted for *DIV*-MARIS indicated for each fly line.

A

B

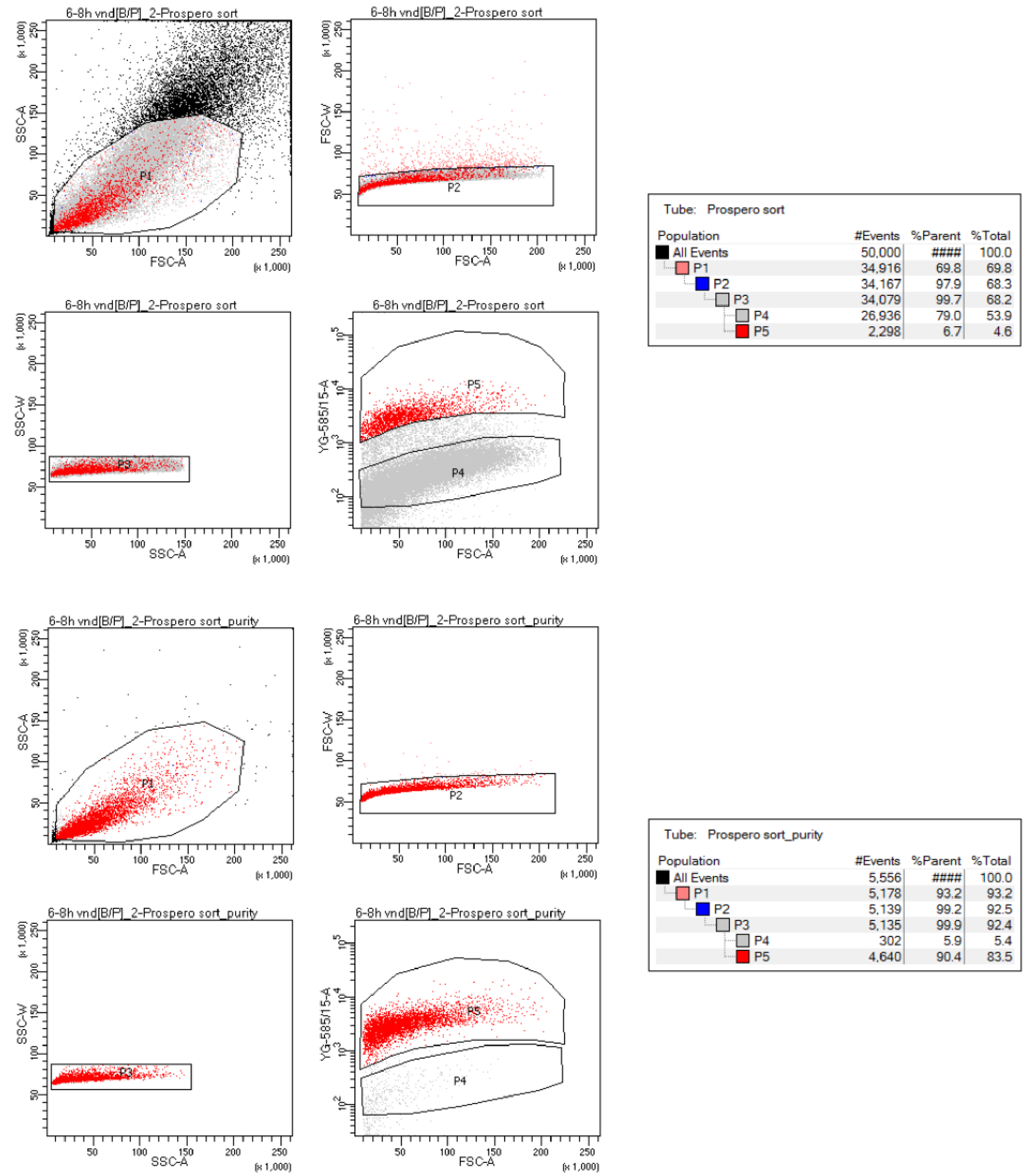

Tube: Prospero sort

| Population | #Events | %Parent | %Total |
|---|---|---|---|
| All Events | 50,000 | #### | 100.0 |
| P1 | 34,916 | 69.8 | 69.8 |
| P2 | 34,167 | 97.9 | 68.3 |
| P3 | 34,079 | 99.7 | 68.2 |
| P4 | 26,936 | 79.0 | 53.9 |
| P5 | 2,298 | 6.7 | 4.6 |

Tube: Prospero sort_purity

| Population | #Events | %Parent | %Total |
|---|---|---|---|
| All Events | 5,556 | #### | 100.0 |
| P1 | 5,178 | 93.2 | 93.2 |
| P2 | 5,139 | 99.2 | 92.5 |
| P3 | 5,135 | 99.9 | 92.4 |
| P4 | 302 | 5.9 | 5.4 |
| P5 | 4,640 | 90.4 | 83.5 |

**Fig. S3: FACS gating strategy yields high purity of sorted populations**
(A) Example FACS gating strategy for sorting of marker-positive (P5; 6.7%) and marker-negative cells (P4; 79%). (B) Re-sort of marker-positive sorted cells shows enrichment of marker-positive (P5; 90.4%) and depletion of marker-negative cells (P4; 5.9%).

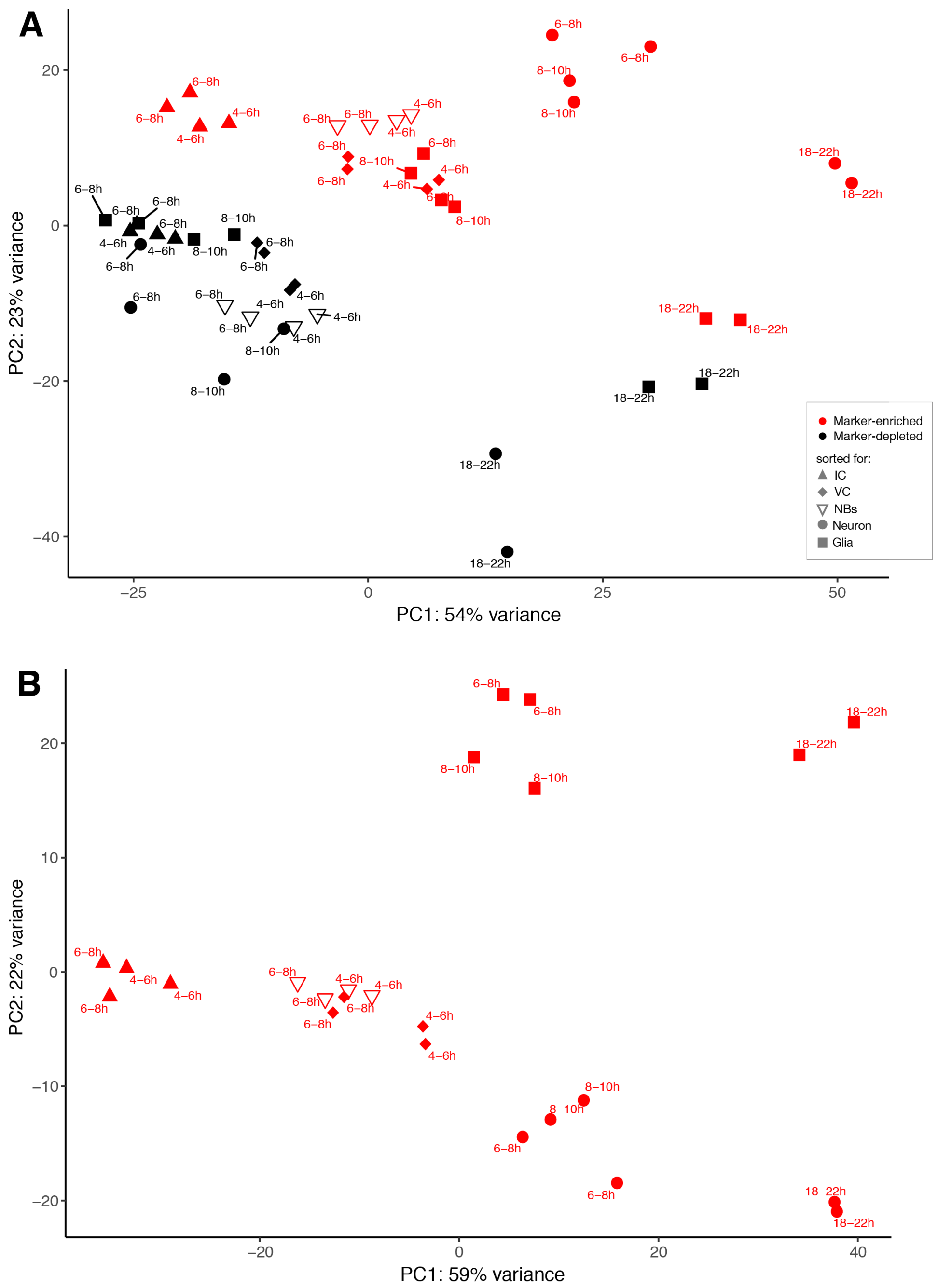
A
PC1: 54% variance
PC2: 23% variance
Marker-enriched
Marker-depleted
sorted for:
IC
VC
NBs
Neuron
Glia
B
PC1: 59% variance
PC2: 22% variance

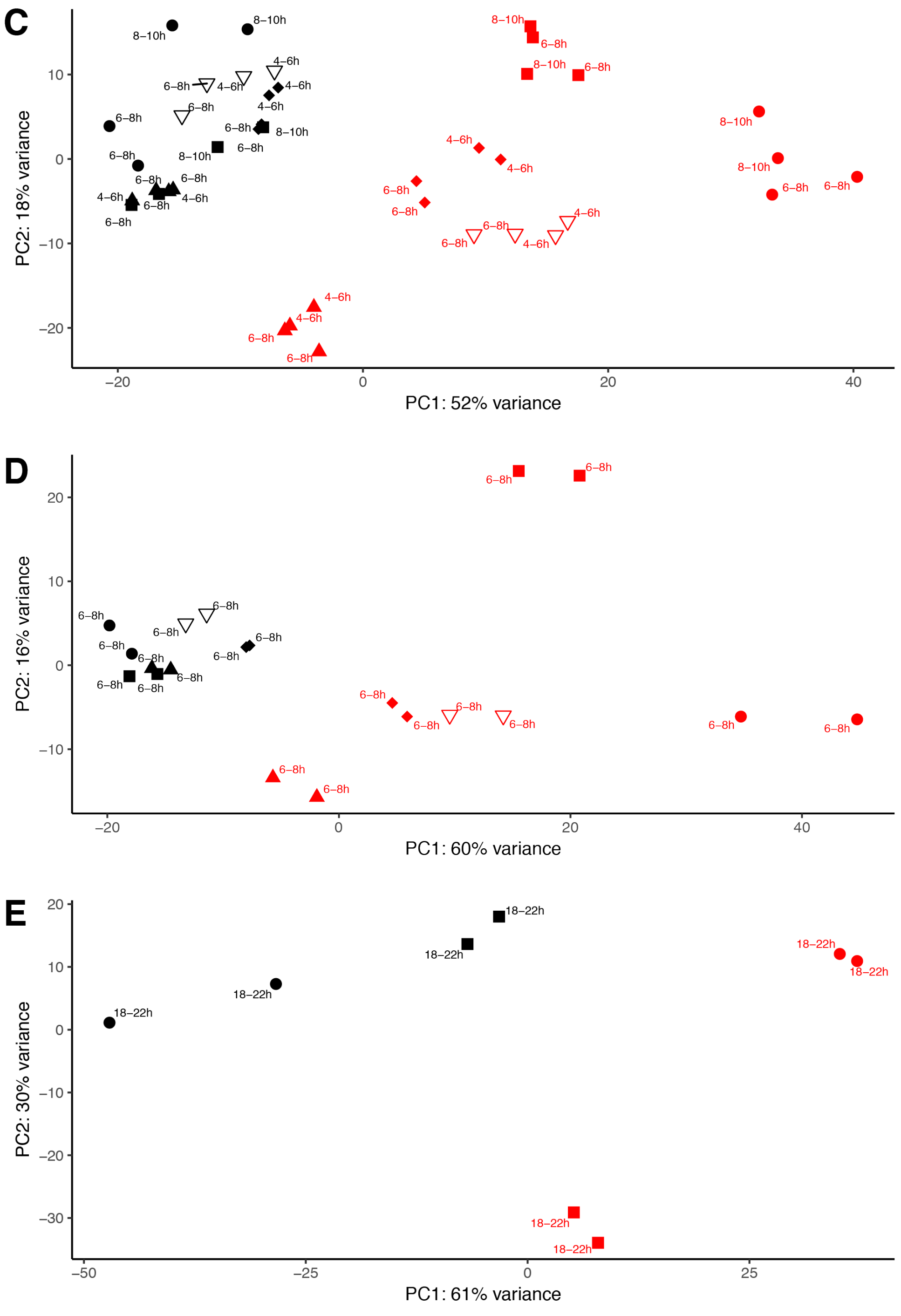
C
PC2: 18% variance
PC1: 52% variance
D
PC2: 16% variance
PC1: 60% variance
E
PC2: 30% variance
PC1: 61% variance

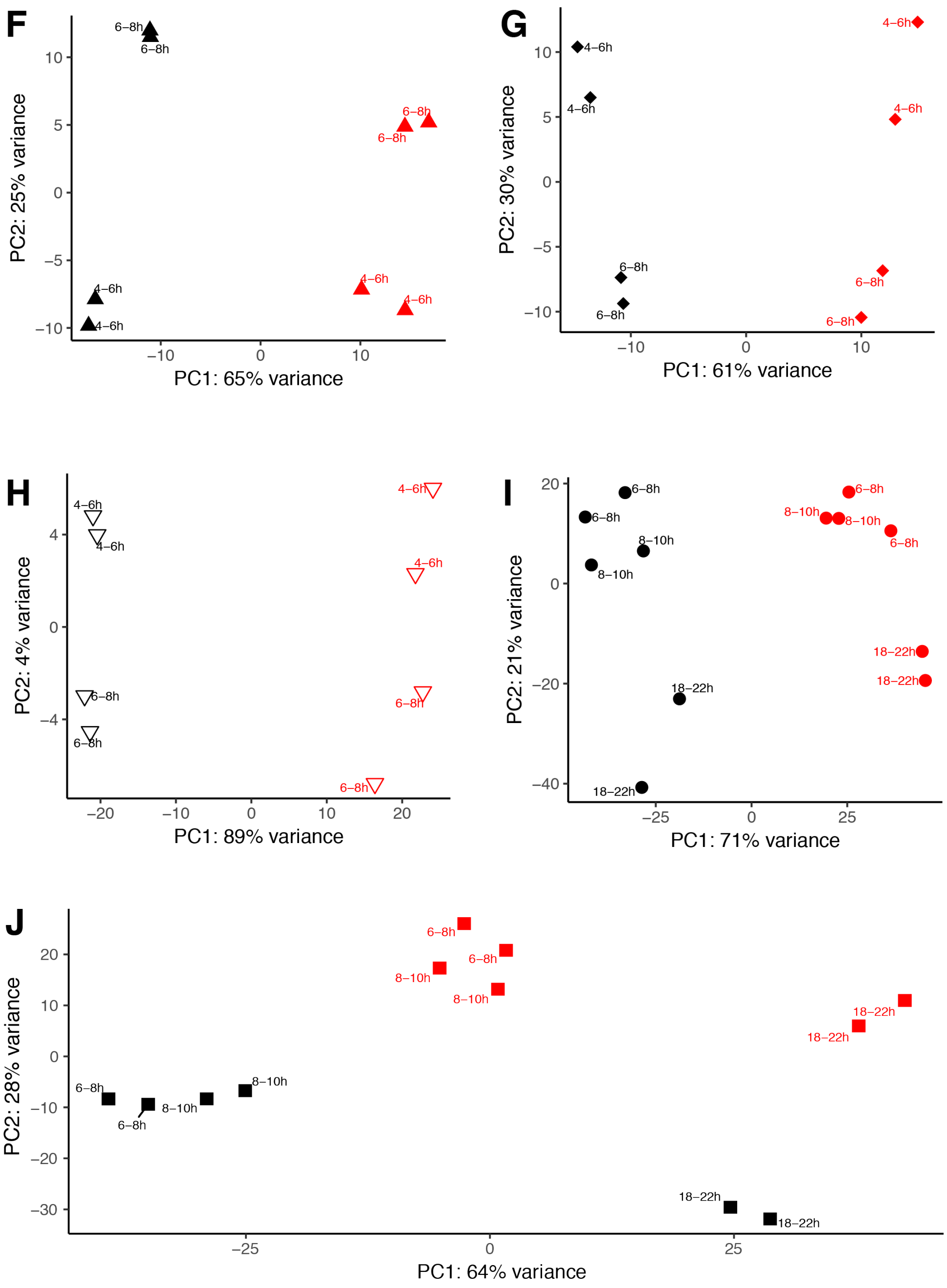

**Fig. S4: Principal component analyses separate spatiotemporal transcriptomes according to tissue and developmental age**

Principal Component Analysis (PCA) with multiple permutations of datasets generated by *DIV*-SortSeq. Marker-enriched datasets depicted in red, marker-depleted datasets in black. (A) All datasets. (B). Only marker-positive datasets. (C). Only early (4-6, 6-8, 8-10h) datasets. (D) Only 6-8h datasets. (E). Only 18-22h datasets. (F). Only datasets sorted for *ind* expression. (G). All datasets sorted for *vnd* expression. (H) All datasets sorted for *pros* expression. (I) All datasets sorted for *elav* expression. (J) All datasets sorted on *repo* expression.

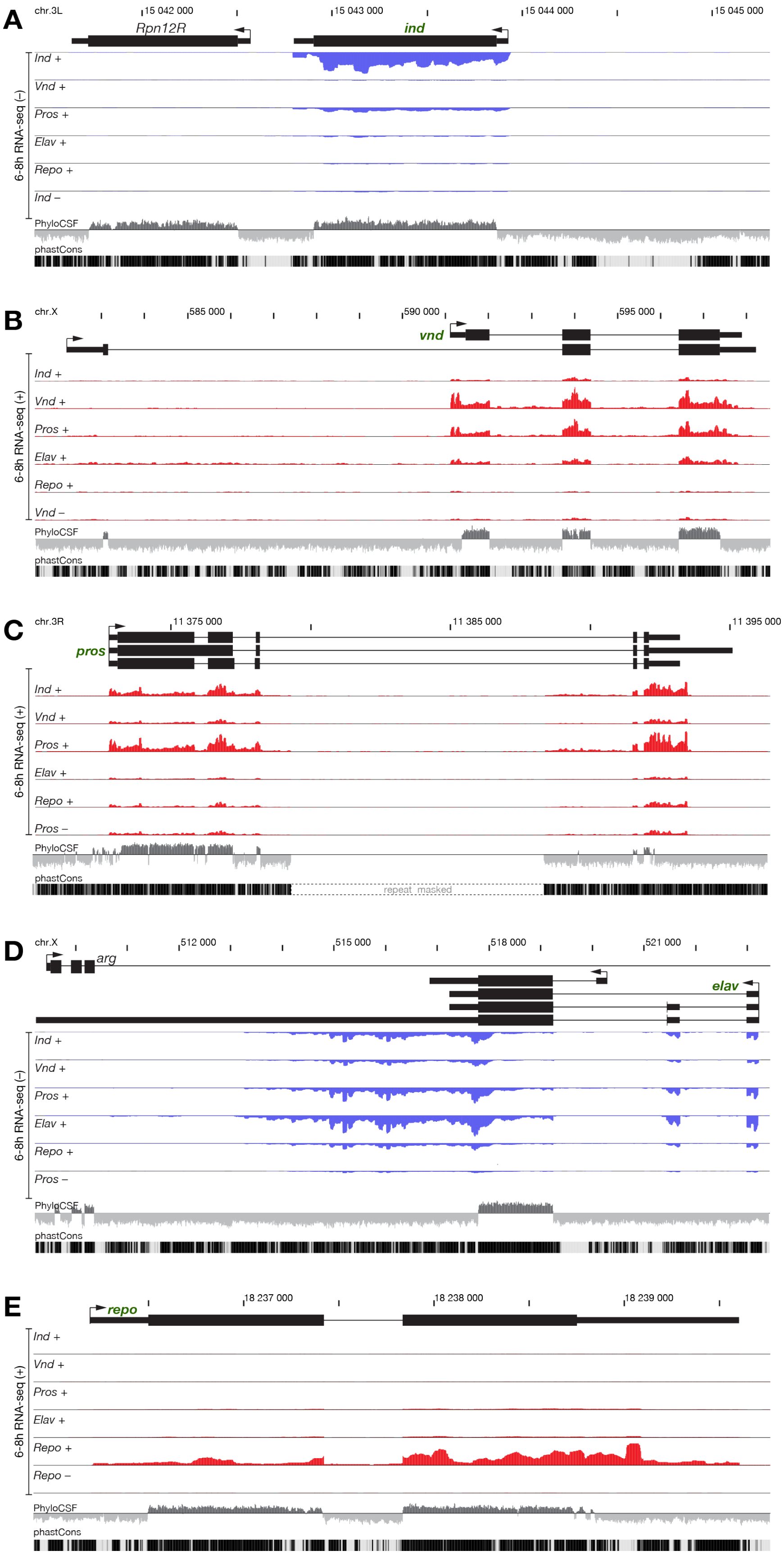
A
chr.3L
15 042 000
15 043 000
15 044 000
15 045 000
Rpn12R
ind
6-8h RNA-seq (–)
Ind +
Vnd +
Pros +
Elav +
Repo +
Ind –
PhyloCSF
phastCons
B
chr.X
585 000
590 000
595 000
vnd
6-8h RNA-seq (+)
Ind +
Vnd +
Pros +
Elav +
Repo +
Vnd –
PhyloCSF
phastCons
C
chr.3R
11 375 000
11 385 000
11 395 000
pros
6-8h RNA-seq (+)
Ind +
Vnd +
Pros +
Elav +
Repo +
Pros –
PhyloCSF
phastCons
repeat masked
D
chr.X
512 000
515 000
518 000
521 000
arg
elav
6-8h RNA-seq (–)
Ind +
Vnd +
Pros +
Elav +
Repo +
Pros –
PhyloCSF
phastCons
E
18 237 000
18 238 000
18 239 000
repo
6-8h RNA-seq (+)
Ind +
Vnd +
Pros +
Elav +
Repo +
Repo –
PhyloCSF
phastCons

**Fig. S5: Cell type specific transcriptomes at neurogenic marker genes**

(A-E) Genome browser data of cell type specific transcriptomes around marker gene loci. Shown are annotated transcripts (top), RNA-seq coverage on the plus- (red) or minus-strand (blue) in the indicated sorts at 6-8hrs AEL, as well as coding potential as measured by PhyloCSF scores (all frames overlaid), and conservation amongst *Drosophilids* (phastCons). (A) *ind* locus. (B) *vnd* locus. (C) *pros* locus. (D) *elav* locus. (E) *repo* locus.

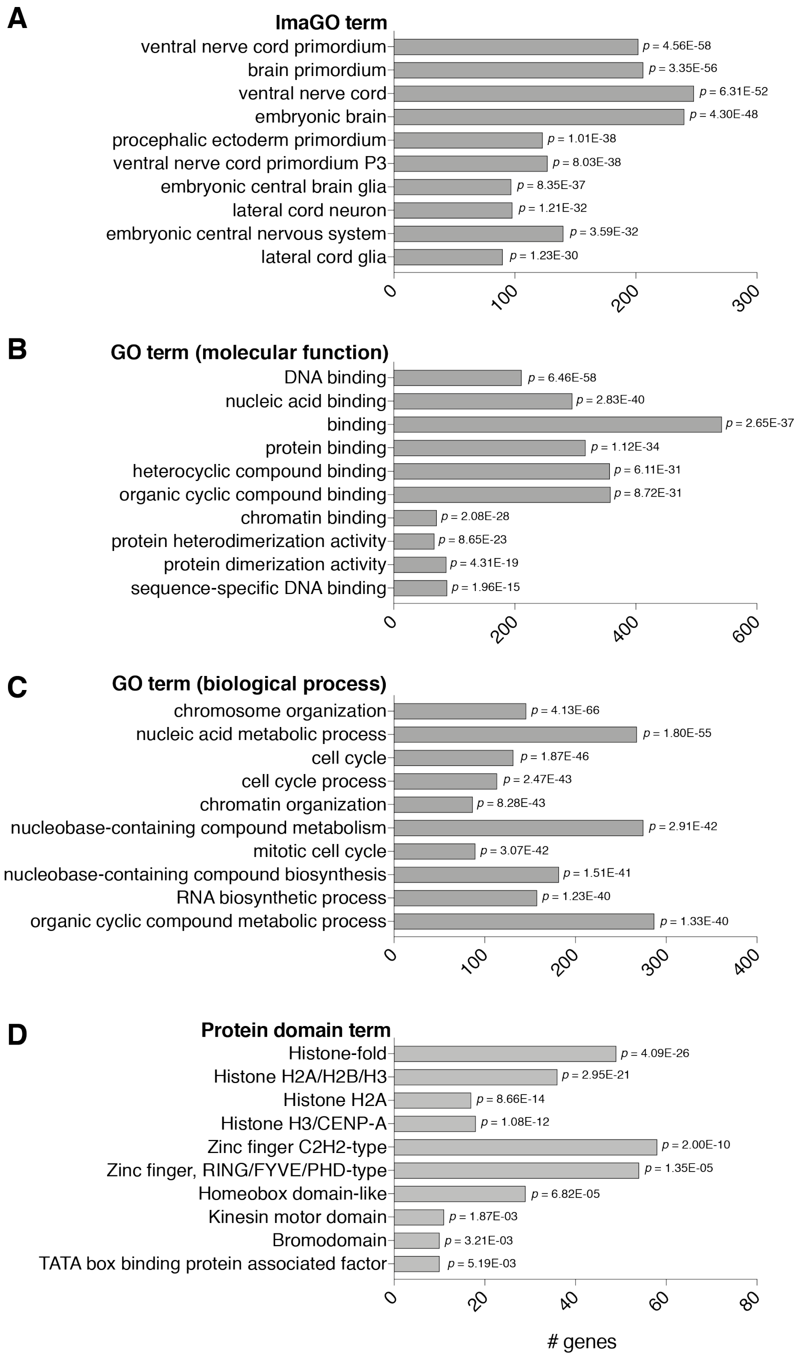

**Fig. S6: Annotations of regulated protein-coding genes predict expression and function**
Analysis of features of 794 protein-coding genes correlated with known neurogenic genes (Pearson correlation, $r > 0.9$) with Flymine {Lyne:2007jd}. (A) Top ImaGO terms. (B) Top GO terms (molecular function). (C) Top GO terms (biological process). (D) Top protein domain terms. Detailed analysis of all genes can be found in File S2.

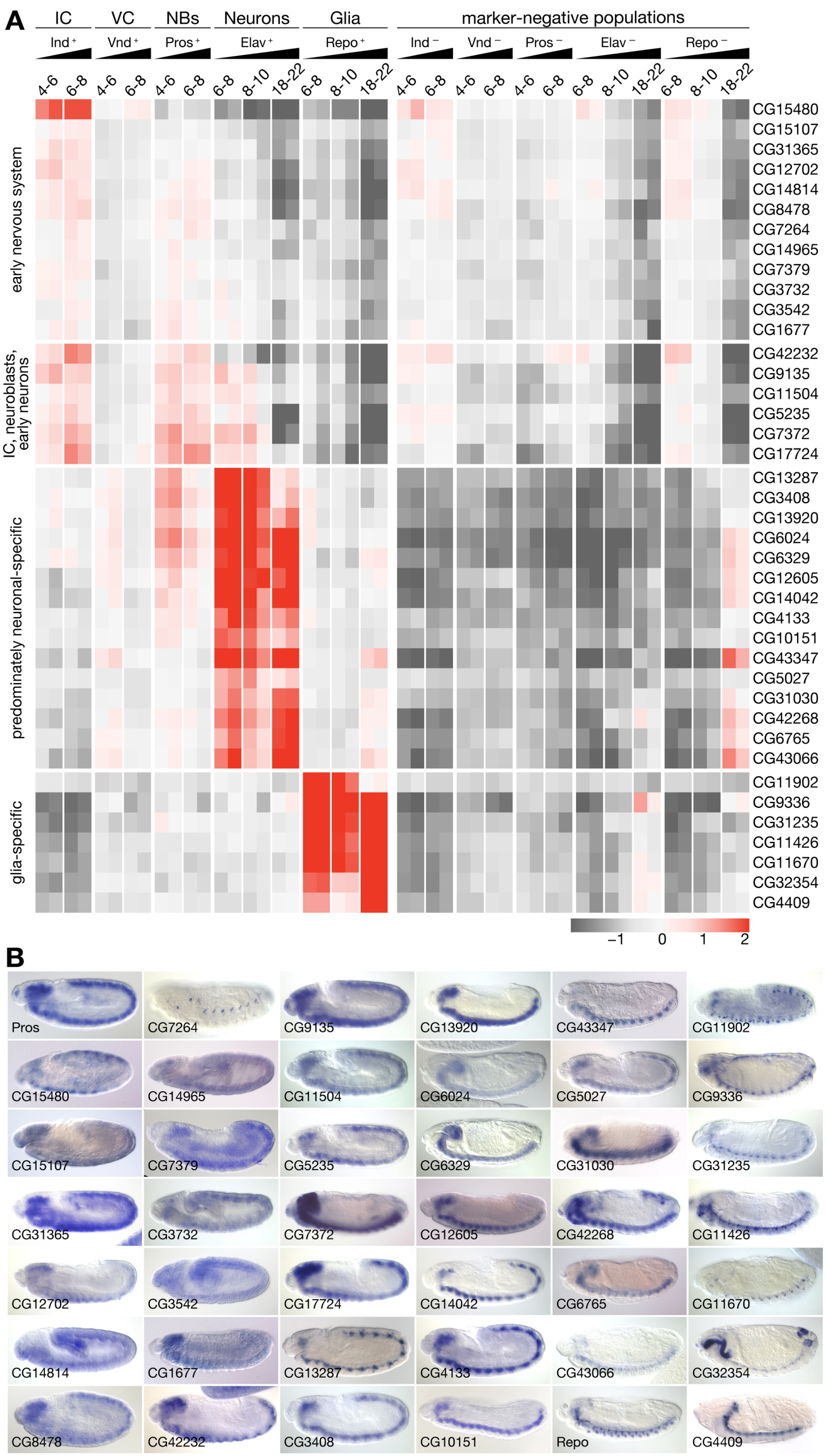
A
IC
VC
NBs
Neurons
Glia
marker-negative populations
Ind +
Vnd +
Pros +
Elav +
Repo +
Ind −
Vnd −
Pros −
Elav −
Repo −
4-6
6-8
8-10
18-22
early nervous system
IC, neuroblasts, early neurons
predominately neuronal-specific
glia-specific
CG15480
CG15107
CG31365
CG12702
CG14814
CG8478
CG7264
CG14965
CG7379
CG3732
CG3542
CG1677
CG42232
CG9135
CG11504
CG5235
CG7372
CG17724
CG13287
CG3408
CG13920
CG6024
CG6329
CG12605
CG14042
CG4133
CG10151
CG43347
CG5027
CG31030
CG42268
CG6765
CG43066
CG11902
CG9336
CG31235
CG11426
CG11670
CG32354
CG4409
−1
0
1
2
B
Pros
CG7264
CG9135
CG13920
CG43347
CG11902
CG15480
CG14965
CG11504
CG6024
CG5027
CG9336
CG15107
CG7379
CG5235
CG6329
CG31030
CG31235
CG31365
CG3732
CG7372
CG12605
CG42268
CG11426
CG12702
CG3542
CG17724
CG14042
CG6765
CG11670
CG14814
CG1677
CG13287
CG4133
CG43066
CG32354
CG8478
CG42232
CG3408
CG10151
Repo
CG4409

**Fig. S7: DIV-SortSeq reveals a multitude of undescribed genes expressed with neurogenic spatiotemporal specificity**

(A) Heatmap of expression profiles of protein-coding genes with unknown function enriched similarly as at least one marker in Fig. 2B (Pearson correlation, $r > 0.9$). Row mean-centered expression values calculated via variance-stabilizing transformation of gene-level RNA-seq counts (scale = $\log_2$ ratio to row mean). (B) Colorimetric RNA *in situ* hybridization of transcripts of unknown function identified in Fig. S7A. All embryos are shown in lateral orientation. Images obtained from Berkeley *Drosophila* Genome Project (Hammonds et al. 2013; Tomancak et al. 2002; Tomancak et al. 2007).

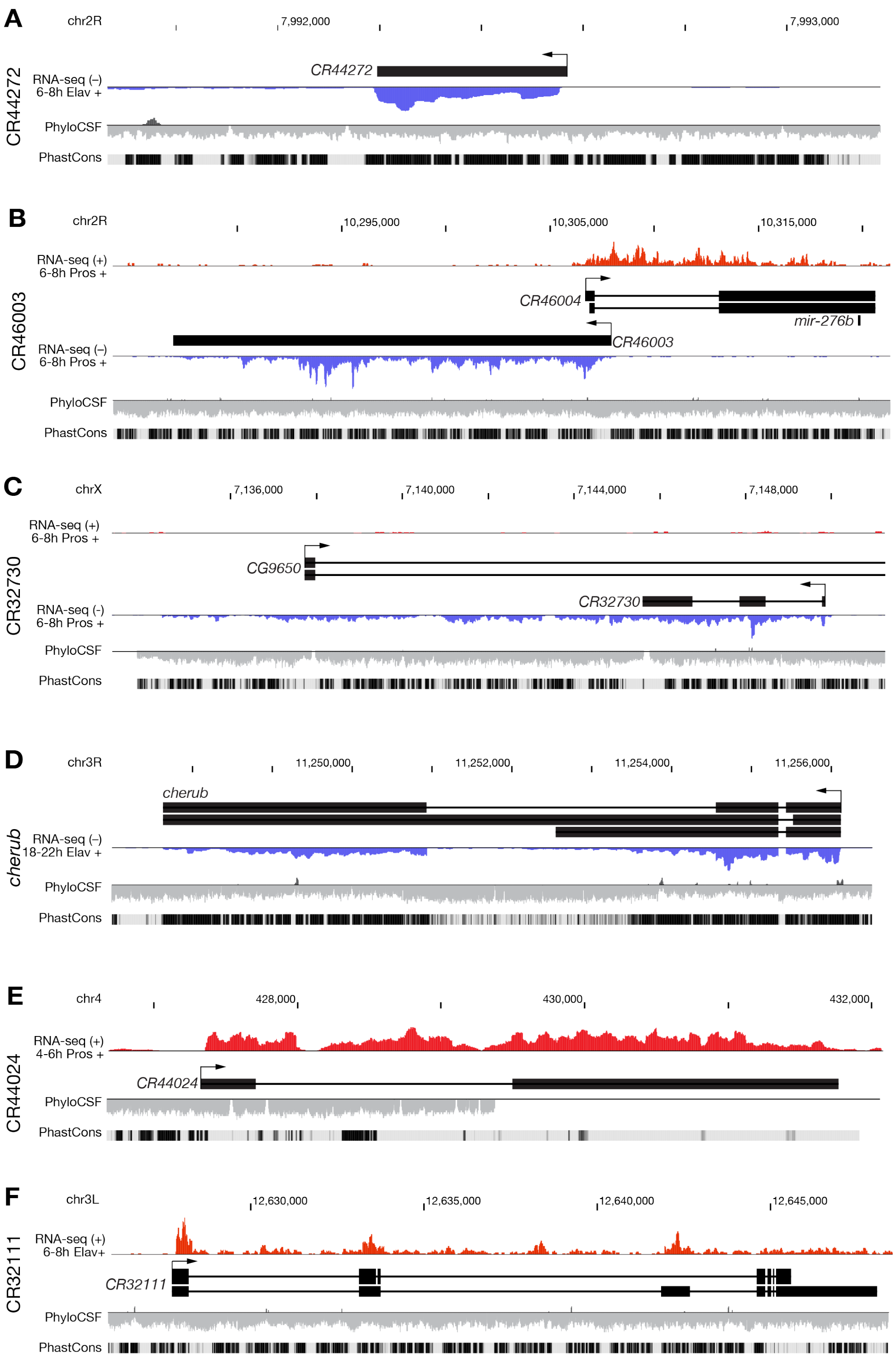
A
chr2R
7,992,000
7,993,000
CR44272
CR44272
RNA-seq (–)
6-8h Elav +
PhyloCSF
PhastCons
B
chr2R
10,295,000
10,305,000
10,315,000
CR46003
RNA-seq (+)
6-8h Pros +
CR46004
mir-276b
CR46003
RNA-seq (–)
6-8h Pros +
PhyloCSF
PhastCons
C
chrX
7,136,000
7,140,000
7,144,000
7,148,000
CR32730
RNA-seq (+)
6-8h Pros +
CG9650
CR32730
RNA-seq (-)
6-8h Pros +
PhyloCSF
PhastCons
D
chr3R
11,250,000
11,252,000
11,254,000
11,256,000
cherub
cherub
RNA-seq (–)
18-22h Elav +
PhyloCSF
PhastCons
E
chr4
428,000
430,000
432,000
CR44024
RNA-seq (+)
4-6h Pros +
CR44024
PhyloCSF
PhastCons
F
chr3L
12,630,000
12,635,000
12,640,000
12,645,000
CR32111
RNA-seq (+)
6-8h Elav+
CR32111
PhyloCSF
PhastCons

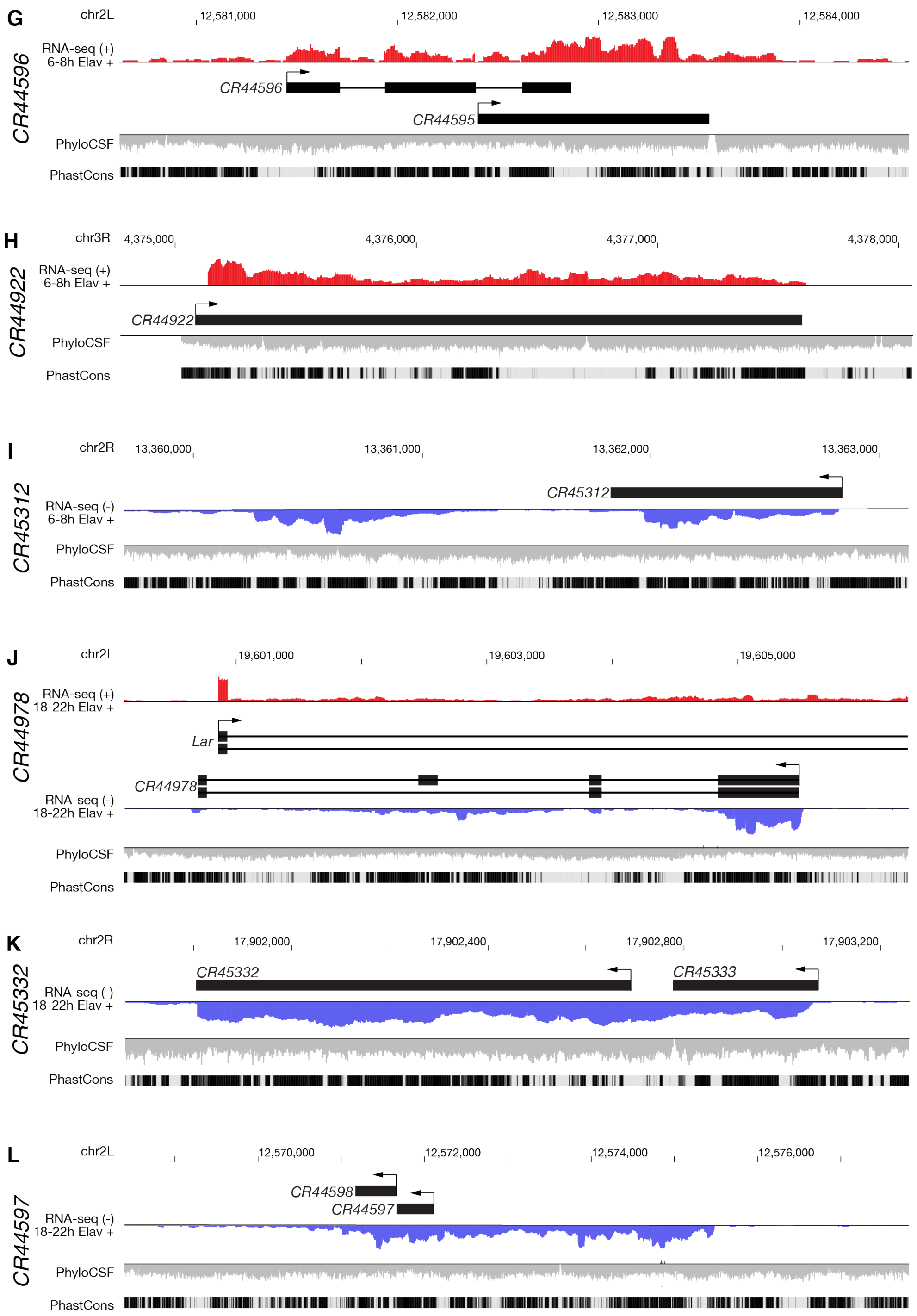

**Figure S8: Cell type specific transcriptomes at neurogenic lncRNA loci**
Browser images of selected lncRNA loci, display as in Figure S4: The (A) *CR44272,* (B) *CR46003* and *CR46004* (pri-miR-276b), (C) *CR32730,* (D) *cherub,* (E) *CR44024,* (F) *CR32111,* (G) *CR44596,* (H) *CR44922,* (I) *CR45312,* (J) *CR44978,* (K) *CR45332* (This gene model might be misannotated – CR45332 and CR45333 might be a single transcript*),* and the (L) *CR44596* and *CR44597* locus.

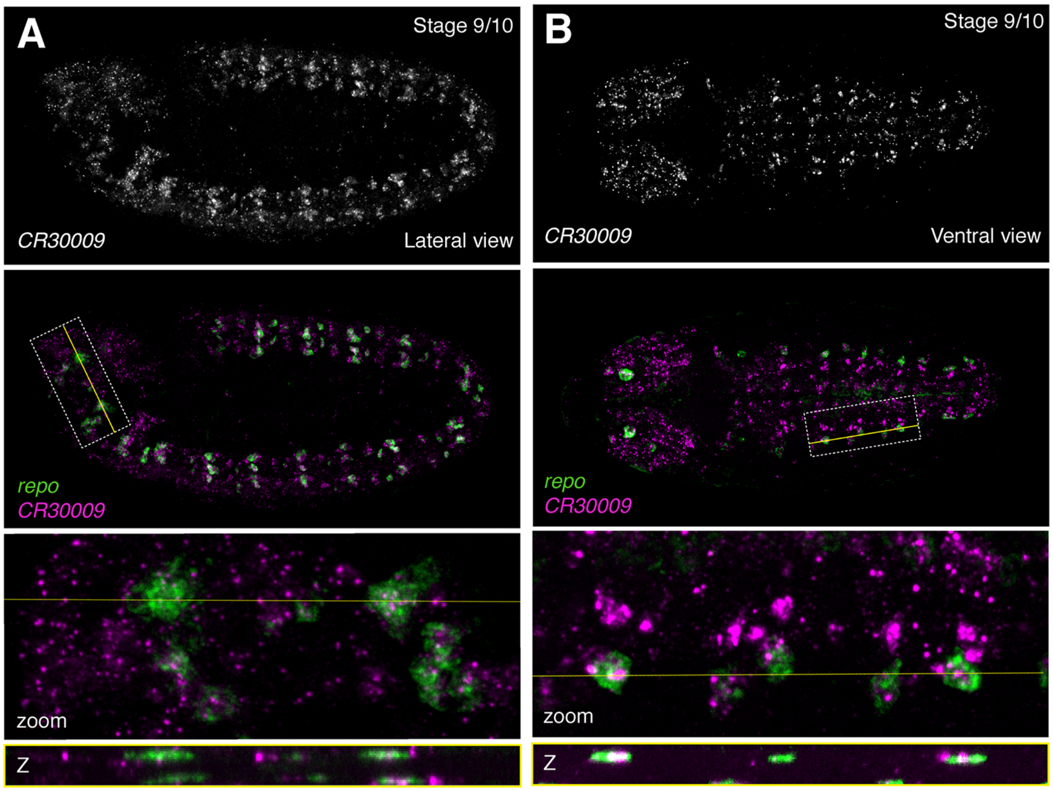

**Fig. S10: The lncRNA *CR30009* is expressed in glial subsets in early embryos**
RNA fluorescent in situ hybridization (RNA-FISH) against *CR30009* and the glial marker *repo*.
(A) Lateral view, stage 9/10. (B) Ventral view, stage 9/10. Top: *CR30009* alone. Second from top: *CR30009* (magenta) overlaid with *repo* (green). Dashed white box indicates region of interest (ROI) and yellow line indicates Z-slice through ROI. Second from bottom: zoom-in of ROI. Bottom: Slice through Z-stack as indicated by yellow line.

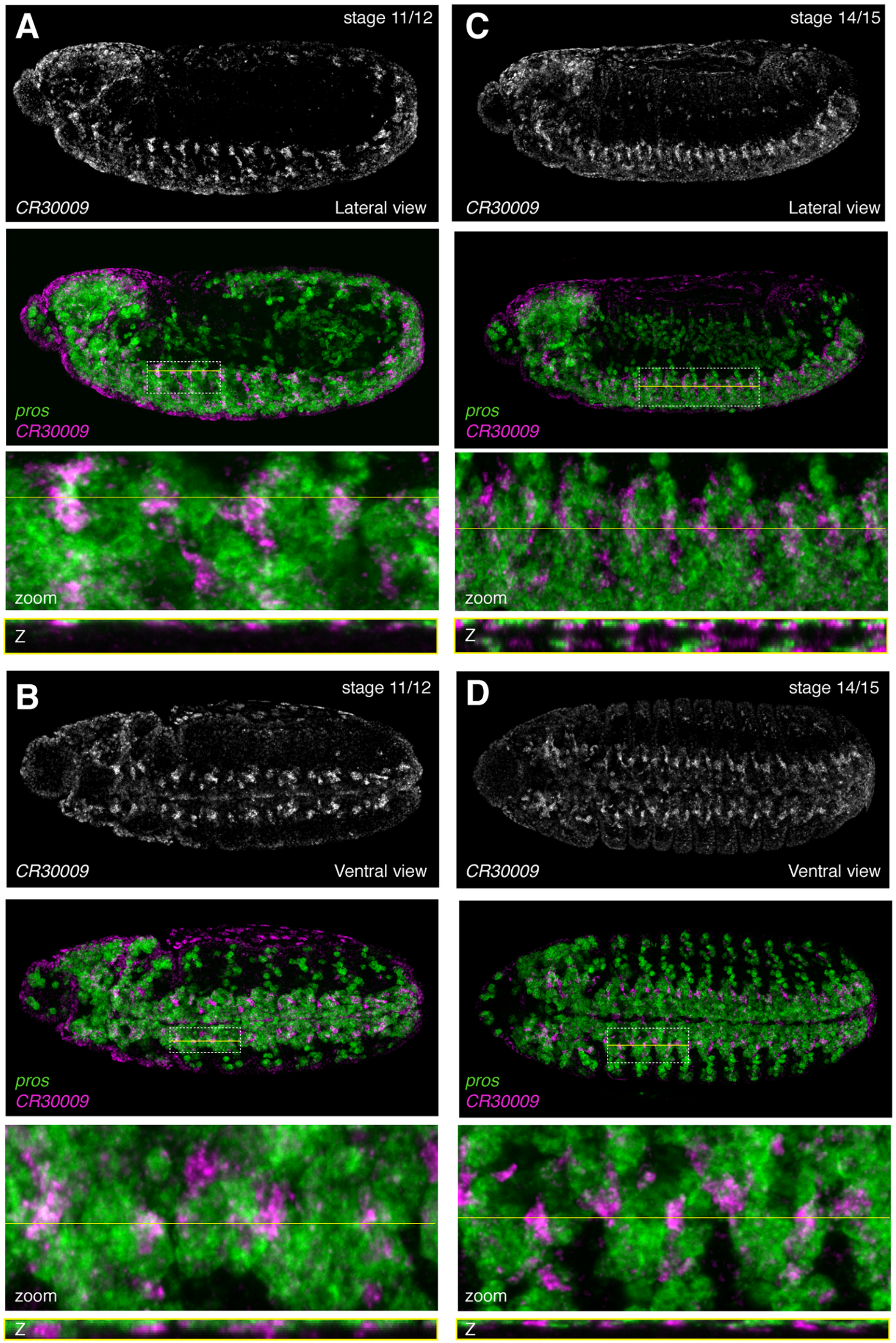

**Fig. S11: The lncRNA *CR30009* is expressed in neuroblast subsets**
RNA fluorescent in situ hybridization (RNA-FISH) against *CR30009* and the neuroblast marker *pros*. (A) Lateral view, stage 11/12. (B) Lateral view, stage 14/15. (C) Ventral view; stage 11/12. (D). Ventral view; stage 14/15. Top: *CR30009* alone. Second from top: *CR30009* (magenta) overlaid with *pros* (green). Dashed white box indicates region of interest (ROI) and yellow line indicates Z-slice through ROI. Second from bottom: zom-in of ROI. Bottom: Slice through Z-stack as indicated by yellow line.

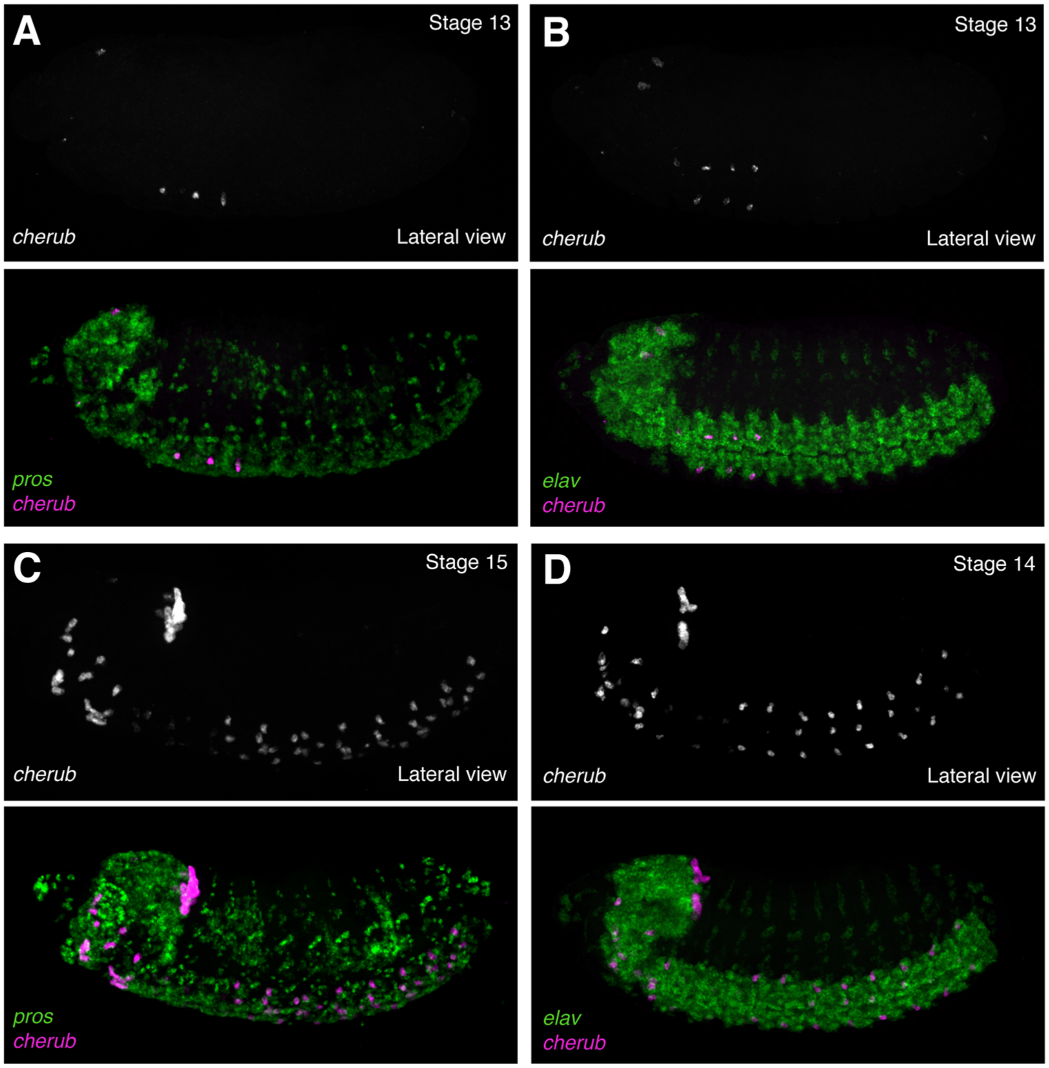

**Fig S12: The lncRNA *cherub* is regulated with strict spatiotemporal specificity.**
RNA fluorescent in situ hybridization (RNA-FISH) against *cherub*, the neuroblast marker *pros*, and the neuronal marker *elav*. Lateral view. (A) *cherub* with *pros*; stage 13. (B) *cherub* with *elav*; stage 13. (C) *cherub* with *pros*; stage 15. (D) *cherub* with *elav*; stage 14. Top: *cherub* alone. Second from top: *cherub* (magenta) overlaid with marker (green).

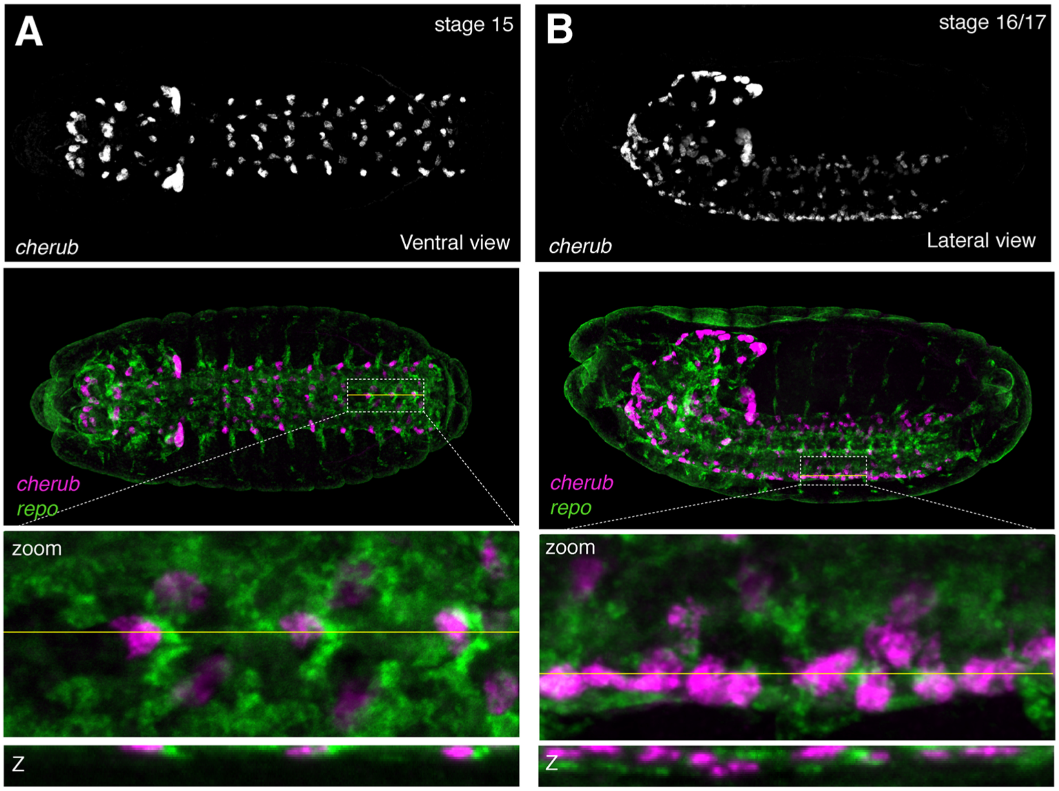

**Fig S13: The lncRNA *cherub* is expressed in glial subsets in late embryos**
RNA fluorescent in situ hybridization (RNA-FISH) against *cherub* and the glial marker *repo*. (A) Ventral view; stage 15. (B) Lateral view; stage 16/17. Top: *cherub* alone. Second from top: *cherub* (magenta) overlaid with *repo* (green). Dashed white box indicates region of interest (ROI) and yellow line indicates Z-slice through ROI. Second from bottom: zom-in of ROI. Bottom: Slice through Z-stack as indicated by yellow line.

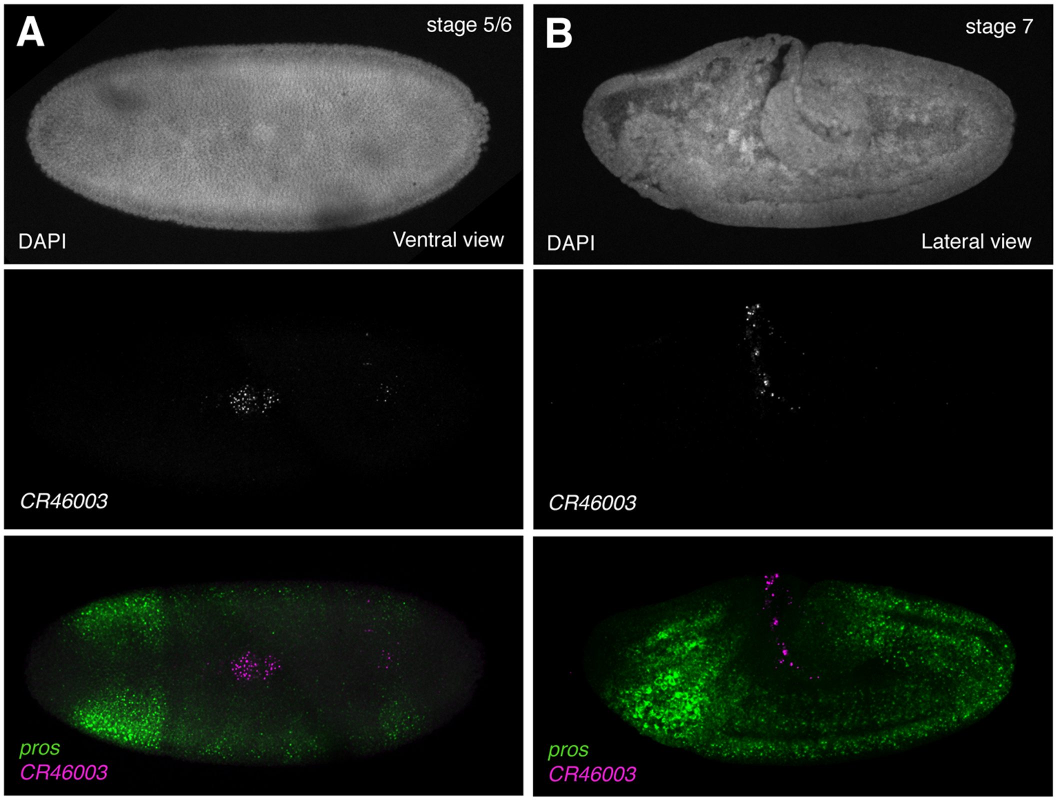

**Fig S14: The lncRNA *CR46003* is expressed in early embryogenesis**
RNA fluorescent *in situ* hybridization (RNA-FISH) against *CR46003* with the neuroblast marker *pros*. (A) Dorsal view; stage 5/6. (B) Lateral view; stage 7/8. Top: DAPI, middle: *CR46003* alone. Bottom: *CR46003* (magenta) overlaid with *pros* (green).

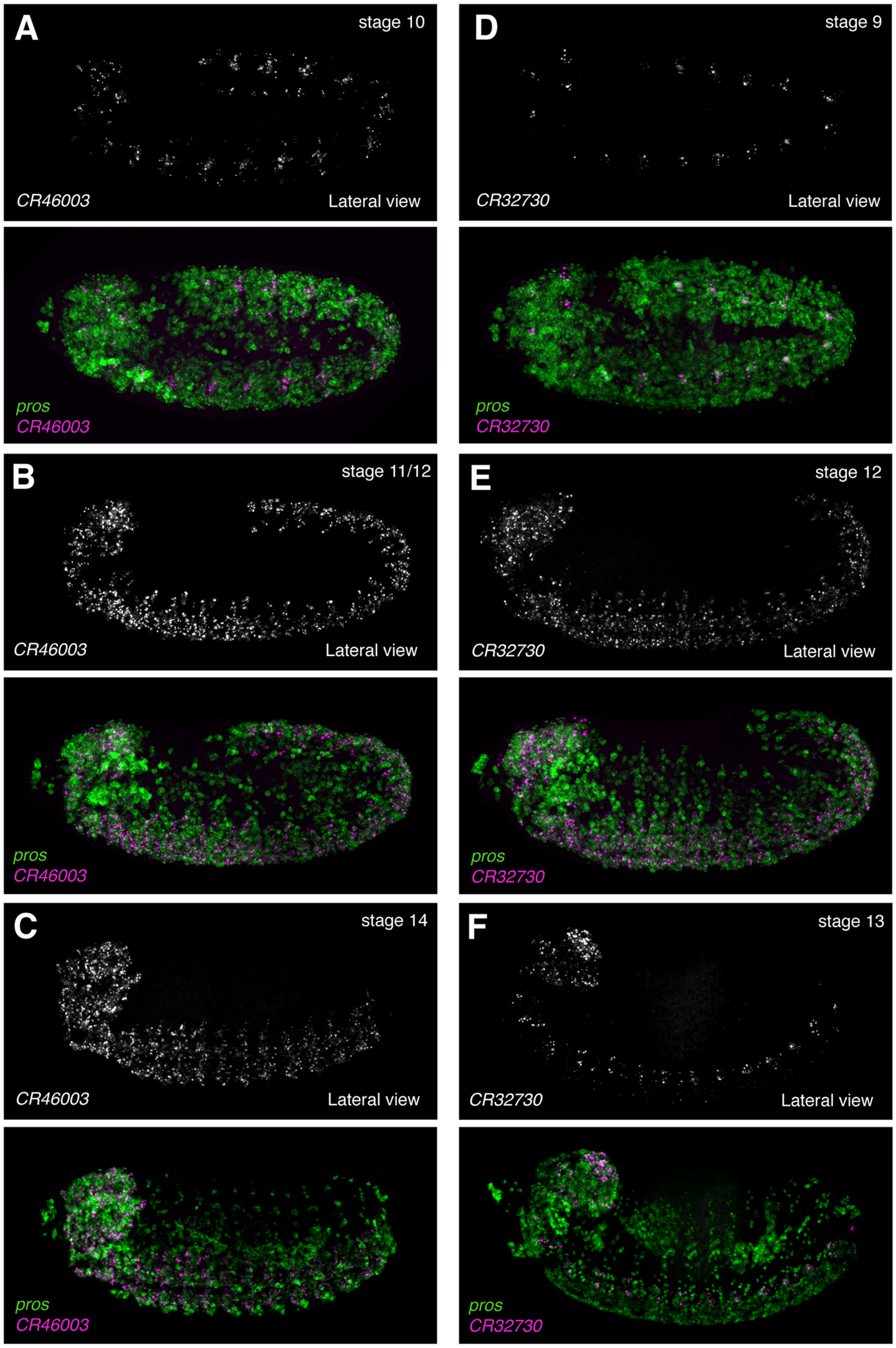

**Fig S15: The lncRNAs *CR46003* and *CR32730* are expressed with similar spatiotemporal specificity.**

RNA fluorescent *in situ* hybridization (RNA-FISH) against *CR46003* and *CR32730* together with the neuroblast marker *pros*. Lateral view. (A) *CR46003*; stage 10. (B) *CR46003*; stage 11/12. (C) *CR46003*; stage 14. (D) *CR32730*; stage 9. (E) *CR32730*; stage 12. (F) *CR32730*; stage 13. Top: lncRNA alone. Second from top: lncRNA (magenta) overlaid with *pros* (green).

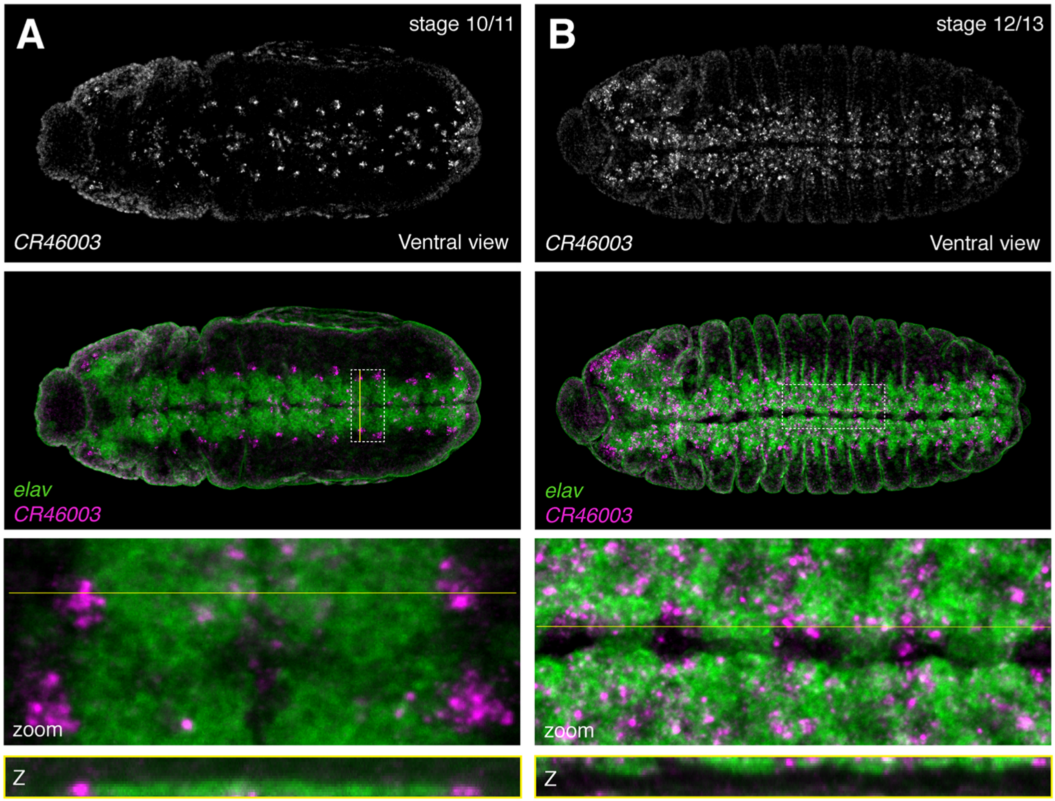

**Fig S16: The lncRNA *CR46003* is expressed in neuronal subsets**

RNA fluorescent in situ hybridization (RNA-FISH) against *CR46003* with the neuronal marker, *elav*. Ventral view. (A) Stage 10/11. (B) Stage 12/13. Top: *CR46003* alone. Second from top: *CR46003* (magenta) overlaid with *elav* (green). Dashed white box indicates region of interest (ROI) and yellow line indicates Z-slice through ROI. Second from bottom: zom-in of ROI. Bottom: Slice through Z-stack as indicated by yellow line.

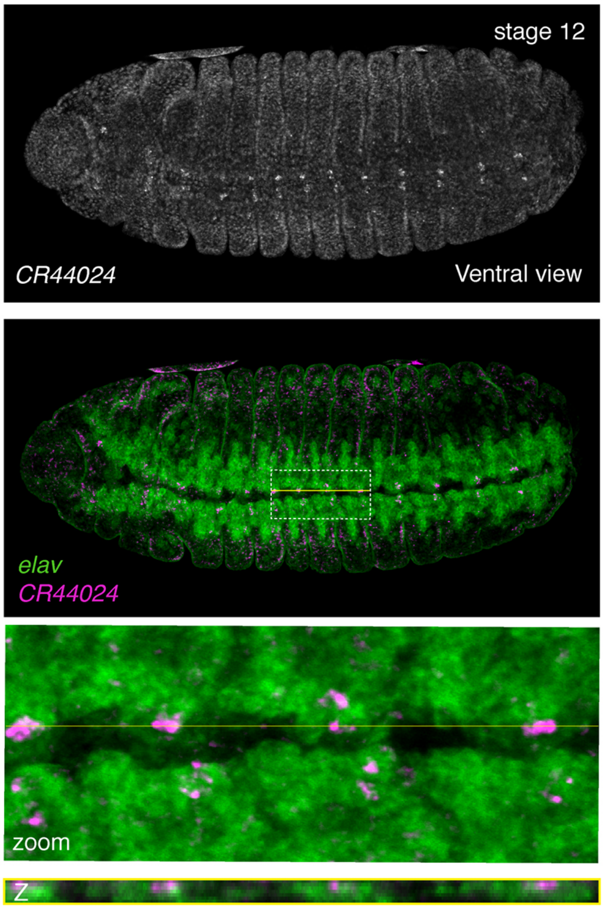

**Fig S17: The lncRNA *CR44024* is expressed from stage 12 in neuronal subsets of the ventral nerve cord.**
RNA fluorescent in situ hybridization (RNA-FISH) against *CR44024* with the neuronal marker, *elav*. Ventral view, stage 12. Top: *CR44024* alone. Second from top: *CR44024* (magenta) overlaid with *elav* (green). Dashed white box indicates region of interest (ROI) and yellow line indicates Z-slice through ROI. Second from bottom: zom-in of ROI. Bottom: Slice through Z-stack as indicated by yellow line.

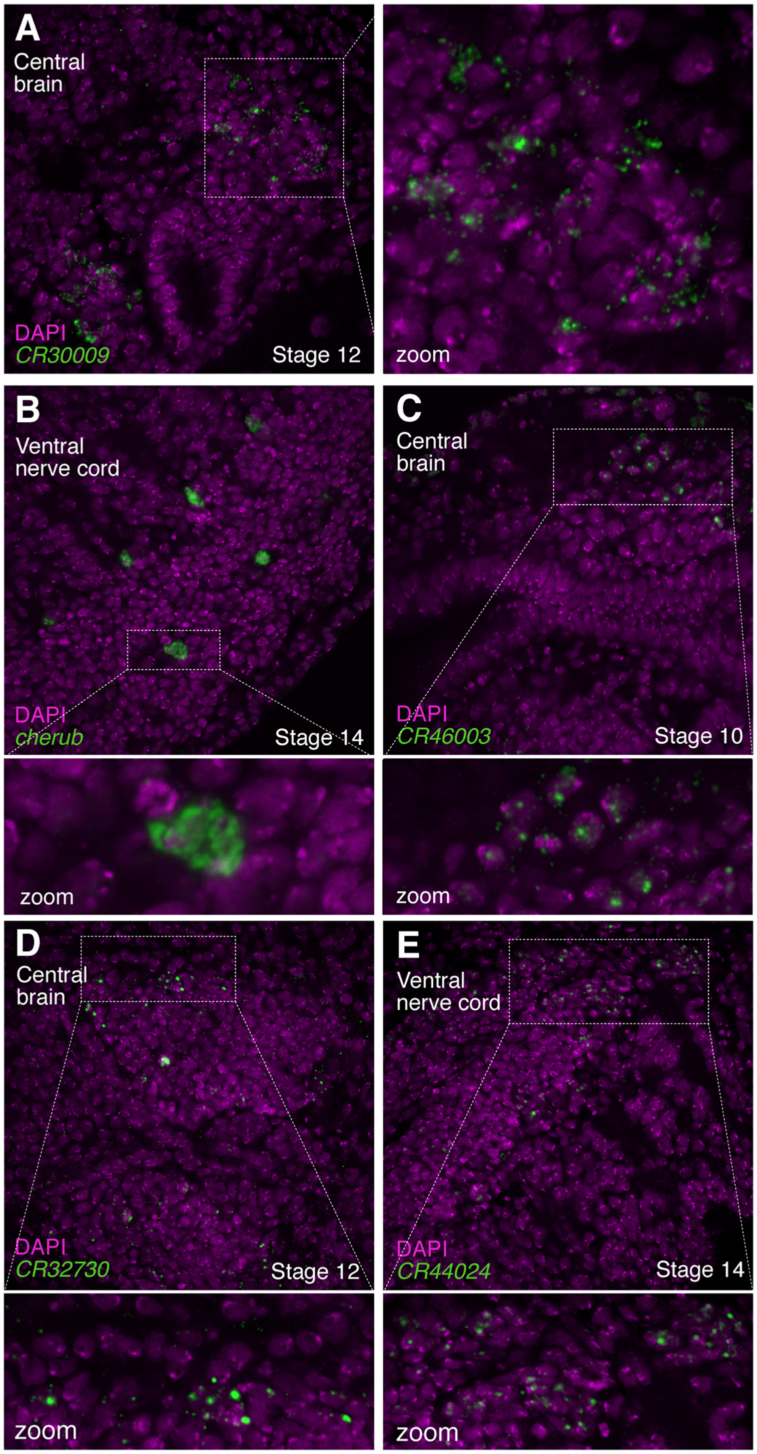

**Fig S18: LncRNAs exhibit varying patterns of subcellular distribution**

RNA fluorescent in situ hybridization (RNA-FISH) against lncRNAs (green) overlaid with the nucleic acid marker, DAPI (magenta). (A) *CR30009*; central brain, stage 12. (B) *cherub*; ventral nerve cord, stage 14. (C) *CR46003*; central brain, stage 10. (D) *CR32730*; central brain, stage 12. (E) *CR44024;* ventral nerve cord, stage 14. (A) Left panel: Dashed white box indicates region of interest (ROI). Right panel: zom-in of ROI. (B-E) Top panel: Dashed white box indicates region of interest (ROI). Bottom panel: zoom-in of ROI.

#### 5. Supplementary References

Barolo, S., Castro, B. & Posakony, J.W., 2004. New Drosophila transgenic reporters: insulated P-element vectors expressing fast-maturing RFP. *BioTechniques*, 36(3), pp.436–40– 442.

Bhatt, D.M. et al., 2012. Transcript dynamics of proinflammatory genes revealed by sequence analysis of subcellular RNA fractions. *Cell*, 150(2), pp.279–290.

Hrvatin, S. et al., 2014. MARIS: Method for Analyzing RNA following Intracellular Sorting K. Aalto-Setala, ed. *PLoS ONE*, 9(3), pp.e89459–6.

Karaiskos, N. et al., 2017. The Drosophila embryo at single-cell transcriptome resolution. *Science*, 358(6360), pp.194–199.

Markstein, M. et al., 2004. A regulatory code for neurogenic gene expression in the Drosophila embryo. *Development*, 131(10), pp.2387–2394.

Pawlicki, J.M. & Steitz, J.A., 2008. Primary microRNA transcript retention at sites of transcription leads to enhanced microRNA production. *The Journal of cell biology*, 182(1), pp.61–76.

Rosner, M. & Hengstschläger, M., 2012. Detection of cytoplasmic and nuclear functions of mTOR by fractionation. *Methods in molecular biology (Clifton, N.J.)*, 821(Chapter 8), pp.105–124.

Rubin, G.M. & Spradling, A.C., 1982. Genetic transformation of Drosophila with transposable element vectors. *Science*, 218(4570), pp.348–353.

Schmittgen, T.D. & Livak, K.J., 2008. Analyzing real-time PCR data by the comparative C(T) method. *Nature Protocols*, 3(6), pp.1101–1108.

Shechner, D.M. et al., 2015. Multiplexable, locus-specific targeting of long RNAs with CRISPR-Display. *Nature Methods*, 12(7), pp.664–670.

Stapleton, M. et al., 2002. The Drosophila gene collection: identification of putative full-length cDNAs for 70% of D. melanogaster genes. *Genome Research*, 12(8), pp.1294–1300.

Vandesompele, J. et al., 2002. Accurate normalization of real-time quantitative RT-PCR data by geometric averaging of multiple internal control genes. *Genome biology*, 3(7), p.RESEARCH0034.
